# Informed, scalable and unified transport couplings for single-cell alignment and downstream inference with scBIOT

**DOI:** 10.64898/2026.01.02.697395

**Authors:** Haihui Zhang, Peiwu Qin

## Abstract

Single-cell genomics requires methods that account for technical variation while preserving biological signals, remain tractable at atlas scale and support analyses beyond visualization, but existing approaches discard the fitted correspondence that could connect these tasks. Here we introduce scBIOT, a biologically informed optimal-transport framework that formulates single-cell alignment as an explicit and inspectable coupling between cells. scBIOT incorporates prior biological knowledge to preserve known cell identities, compresses transport through prototypes for large datasets and quantifies gene-level changes induced by alignment using coupling-derived transport scores. Across transcriptomic and spatial mapping benchmarks, scBIOT balanced biological conservation, batch correction and label-transfer accuracy while distinguishing batch-associated stress responses from biological immune and drug-response programs. Prototype transport scaled coupling-based analysis to approximately 100 million perturbed cells. Because the coupling encodes probabilistic correspondences rather than only corrected coordinates, the same fitted coupling supports state mapping, trajectory inference, perturbation analysis, multimodal comparison and spatiotemporal localization. Applications to cancer, hematopoiesis and embryonic development identified condition-specific cell states, modality-dependent regulatory programs and spatially resolved developmental transitions. Informed, scalable and unified transport couplings therefore link atlas alignment to downstream biological inference within a single framework.

## Introduction

Single-cell omics technologies have transformed the study of cellular heterogeneity, states and dynamics across diverse biological and biomedical contexts. However, integrating datasets across experiments or modalities remains challenging because technical artifacts and modality-specific differences can obscure biological signals and confound downstream analyses. Single-cell atlases are increasingly assembled across individuals, experiments and technologies and used as references onto which new observations are mapped. This strategy makes cell states comparable at a scale inaccessible to isolated studies, but it also creates a fundamental inference problem. Technical variation should be reduced without eliminating biological differences associated with assays or modalities^1–3^. An integration method should therefore do more than mix batches or align modalities. It should use available biological information, scale to atlas level and reveal enough of its fitted transformation for investigators to assess what the alignment preserved or altered.

Existing approaches solve important parts of this problem. Reference-mapping methods project query cells into a fixed atlas, probabilistic models learn shared latent variables and anchor-based procedures identify local correspondences^4–8^. Their performance is generally evaluated through complementary measures of batch removal, biological conservation and label-transfer accuracy^1^. These measures are indispensable for comparing embeddings, but they do not identify the genes or programs most altered by a fitted alignment. This limitation becomes consequential when a biological axis is partially confounded with batch or modality, because a visually well-mixed representation can conceal loss of the signal that motivated the experiment and compromise label transfer and downstream analyses.

The same fragmentation occurs after single-cell alignment. Differential-state analysis, multimodal comparison, perturbation routing and temporal or spatial mapping are commonly implemented as separate workflows, each with its own representation and correspondence model. This separation makes it difficult to relate downstream biological findings to the alignment that generated them. It can also discard information, whereas an optimal transport (OT) fit retains not only mapped coordinates but also an explicit coupling that records how mass is distributed across cellular states. OT provides a principled, geometry-aware way to compare probability distributions and has already proved useful across single-cell tasks, including temporal mapping^4,9^, trajectory inference^10–12^, spatial alignment^13,14^, spatial cell-cell communication^15^, spatial gene reconstruction^16,17^, cross-modality integration^14,18^ and perturbation analysis^19,20^. However, atlas-integration workflows generally reveal little about the molecular basis of the inferred correspondence. scBIOT addresses this gap through a coupling-conditioned gene-level audit that projects reference expression through the inferred correspondence and ranks genes and programs that remain discordant between aligned states. Together with biologically informed transport and density-weighted prototypes, this framework makes atlas integration inspectable at gene resolution, scalable to atlas-sized datasets and applicable across downstream multiomic analyses.

We developed scBIOT to make this coupling the central analytical object. scBIOT learns a compact expression geometry, constructs an entropically regularized unbalanced OT coupling between domains and retains the coupling for subsequent analyses. Known cell identities can shape the shared embedding and constrain integration in supervised scBIOT (supBIOT), reducing biologically implausible matches during reference mapping. Row-normalized transport weights drive barycentric mapping, and the resulting aligned embedding supports label transfer. The same weights can be projected back to gene space to quantify which molecular features were moved, aggregated across states to compare biological conditions, or composed across modalities, perturbations and developmental stages. For very large datasets, density-weighted prototypes preserve state-level structure while avoiding intractable cell-to-cell coupling.

We evaluated the framework in a sequence designed to test its three claims. First, heterogeneous transcriptomic and spatial benchmarks tested whether biologically informed transport improved mapping without sacrificing batch correction. Second, deliberately controlled analyses asked whether the retained coupling made integration auditable at gene resolution and whether prototype transport remained usable at the scale of 100 million cells. Third, we applied a shared coupling-based framework to investigate immune states in head and neck squamous cell carcinoma (HNSCC), coordinated and modality-biased programs in multiomics, and spatially localized developmental transitions in mouse organogenesis. Across these settings, scBIOT linked alignment to interpretation while preserving a clear distinction between inferred correspondence, biological replication and causal evidence.

## Results

### scBIOT links alignment and downstream inference through the fitted transport coupling

scBIOT was designed to address a limitation of integration workflows that typically retain only the corrected representation while discarding the fitted correspondence that generated it. By retaining the transport coupling, scBIOT links alignment directly to downstream inference and preserves evidence of how the corrected representation was obtained. The framework first uses a tied linear encoder and decoder to learn a compact representation of gene expression (Fig. 1a). The orthogonality-constrained linear map discourages redundant latent axes while maintaining a direct connection to gene space. OT is then solved between cells or density-weighted prototypes in this representation rather than in the original sparse, high-dimensional matrix.

**Fig. 1.**
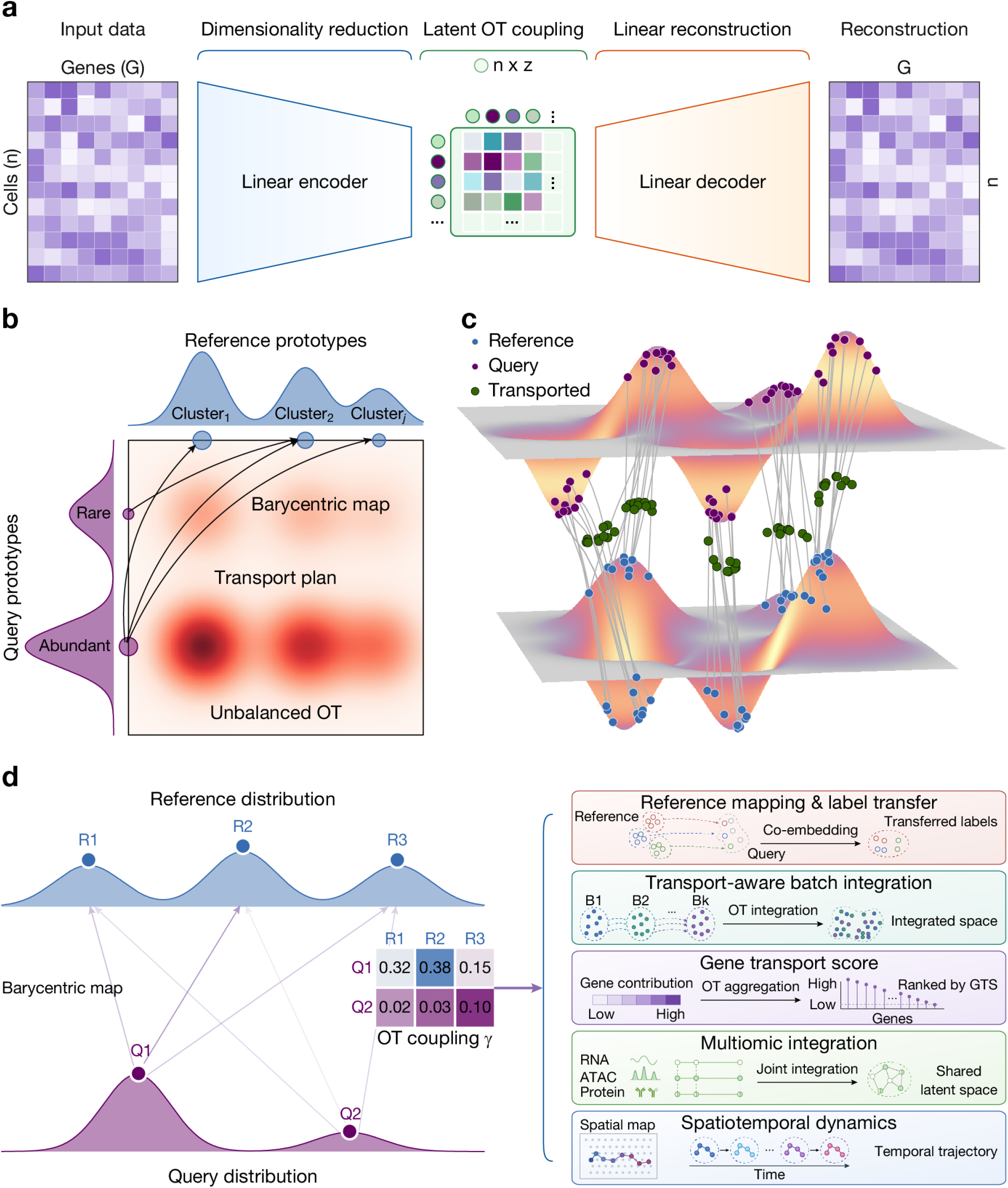
scBIOT retains an explicit optimal-transport coupling for downstream analysis. **a**, A tied linear encoder maps a cell-by-gene matrix into a latent representation, and a tied decoder reconstructs gene space. **b**, Unbalanced prototype OT couples query and reference states while permitting unmatched mass. **c**, The coupling defines barycentric positions between query and reference distributions. **d**, The retained matrix supports reference mapping, batch integration, gene transport scoring, multiomic analysis and spatiotemporal transport.

The coupling is entropically regularized to stabilize estimation and unbalanced to accommodate unequal mass between the two domains (Fig. 1b). This is important when the query contains a state that is absent or substantially depleted in the reference, because balanced transport would distribute all query mass even when no close biological counterpart exists. In the unbalanced formulation, marginal penalties discourage arbitrary mass loss without requiring every prototype to be fully matched. Row normalization yields conditional transport weights, whose coupling-weighted reference coordinates define the barycentric position of each query cell or prototype (Fig. 1c).

Biological information can shape the shared embedding and constrain integration when reference labels are available. In supBIOT, a class-supervised encoder and a label-guided integration step favor compatible cell identities during fitting, which constrains alignment without requiring labels for every query state. The fitted weights are then used for mapping and label transfer. More broadly, scBIOT treats transport couplings as explicit analytical objects rather than disposable intermediates. Projection through the fitted coupling quantifies gene-level displacement, and aggregation of signed displacement compares biological conditions. Cross-modal couplings compare molecular programs, while compositions of couplings estimated between adjacent stages support temporal pullback (Fig. 1d). Figure 1 therefore defines the common computational framework, while the subsequent analyses evaluate its informed, scalable and unified applications.

### Biologically informed transport couples biological conservation with accurate label transfer

We first asked whether constraining transport with known biological identities could improve reference mapping across studies and measurement platforms. We compared supBIOT with Symphony, scANVI, scPoli, Seurat and an unintegrated baseline^5–8^. The benchmark included pancreas and peripheral blood mononuclear cell (PBMC) collections, the cross-tissue Tabula Muris Senis and Tabula Sapiens atlases, and Xenium datasets from colorectal, liver and ovarian cancers^1,21–24^. These tasks differ in species, tissue complexity, class balance and assay geometry, providing a heterogeneous evaluation rather than repeated partitions of a single benchmark.

Because no single metric captures successful mapping, we summarized performance across datasets using the total scIB integration score, which combines biological conservation and batch correction, together with weighted and macro label-transfer F1 scores. supBIOT achieved the highest mean integration score (0.680), compared with 0.658 for the second-ranked method, scANVI (Fig. 2a), indicating that supervision did not simply trade batch correction for class separation. supBIOT also achieved the highest mean weighted F1 (0.878), exceeding the second-ranked method, Seurat (0.829; Fig. 2b), and the highest mean macro F1 (0.711), exceeding the second-ranked method, Symphony (0.621; Fig. 2c). These corresponded to gains of 4.8 percentage points in weighted F1 and 9.0 percentage points in macro F1 over the respective runners-up. All methods were evaluated on identical query cells and ground-truth labels using the same scoring procedure within each dataset, so these comparisons reflect differences between methods rather than preprocessing or evaluation. supBIOT additionally ranked first for both F1 measures in each of the seven benchmarks. Thus, biologically informed transport provided the strongest average integration and label-transfer performance across metrics that emphasize both abundant and low-frequency cell classes.

**Fig. 2.**
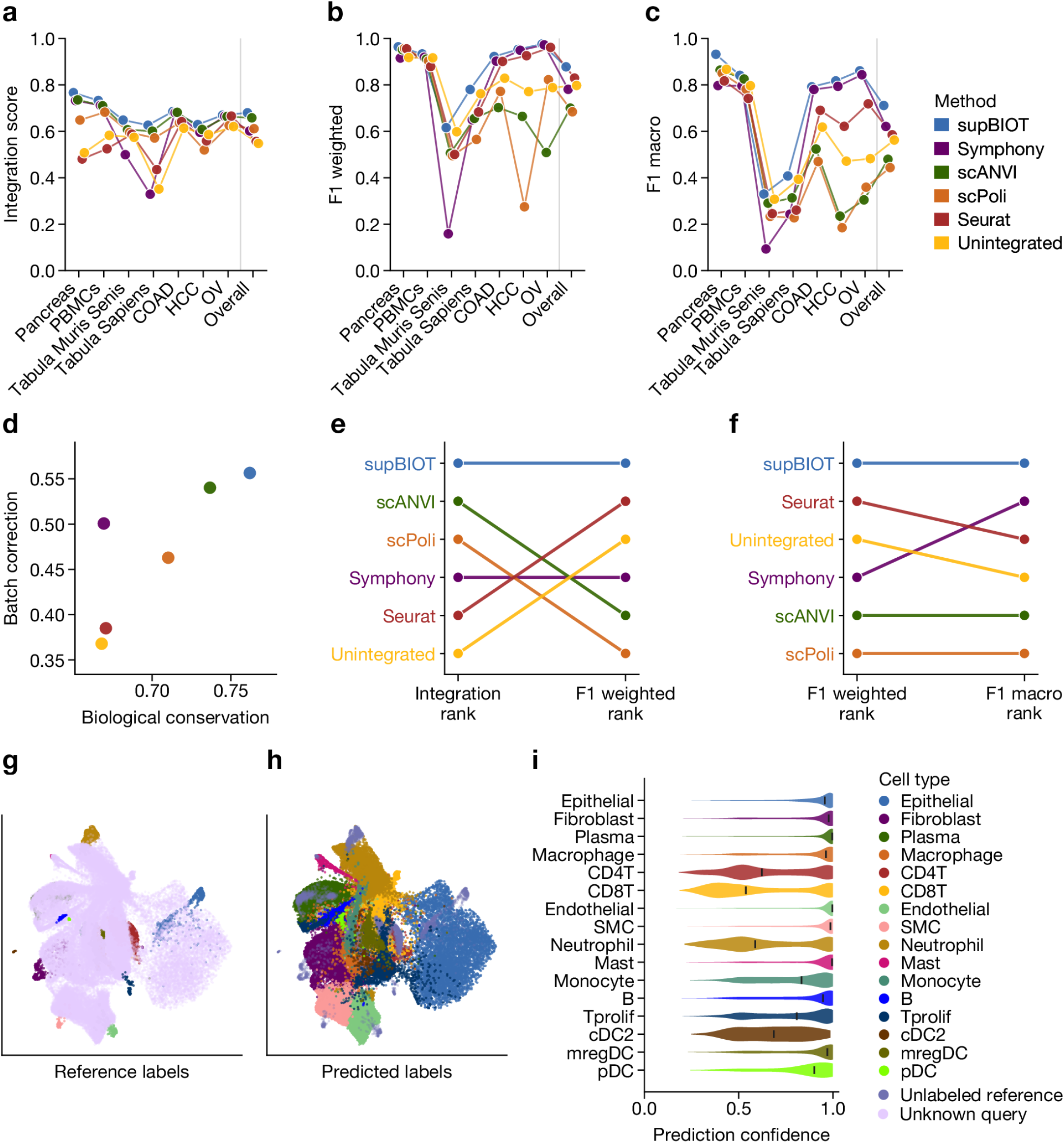
Reference mapping and label transfer across heterogeneous datasets. The compared methods are supBIOT, Symphony, scANVI, scPoli, Seurat and unintegrated PCA. **a**, Total integration score across pancreas, PBMC, Tabula Muris Senis, Tabula Sapiens and three Xenium datasets. **b**, Weighted label-transfer F1. **c**, Macro label-transfer F1. Dots connected within a method represent datasets; “Overall” denotes the across-dataset summary. All methods were scored on identical query cells, labels and metric code within each dataset. **d**, Mean biological-conservation and batch-correction scores. **e**, Rank changes from the integration score to weighted F1. **f**, Rank changes from weighted to macro F1. **g**,**h**, Xenium colorectal-cancer query colored by reference and transferred labels. **i**, Per-cell prediction confidence by predicted cell type.

Decomposition of the integration score clarified the basis of this performance. In the across-dataset summary, supBIOT occupied the region of high biological conservation while retaining substantial batch correction (Fig. 2d). This pattern is consistent with the intended role of informed transport, which uses biological information to restrict implausible correspondences rather than maximizing mixing irrespective of cell identity. Method rankings nevertheless changed between the total integration score and weighted F1 (Fig. 2e), and again between weighted and macro F1 (Fig. 2f). Thus, an embedding that performs well globally may not provide equally reliable annotation, and a method that annotates abundant populations well may not do so for low-frequency ones. In contrast to the rank shifts observed for competing methods, supBIOT remained the top-ranked method for integration, weighted F1 and macro F1, indicating consistent performance across global alignment and label-transfer criteria.

We next examined the fitted mapping at single-cell resolution using a representative Xenium colorectal cancer query. Reference annotations and predicted labels showed coherent organization in the shared representation (Fig. 2g,h), whereas prediction-confidence distributions differed across cell types (Fig. 2i). These cell-type-specific distributions exposed uncertain mappings that were not apparent from the global benchmark scores and could therefore guide manual inspection. Together, these results show that the supervised scBIOT formulation uses biological information to preserve cell-identity structure while correcting cross-dataset variation, establishing the first sense in which the method is informed.

### Transport scores quantify integration-induced molecular displacement

To inform the molecular interpretation of integration, we sought to quantify which genes and biological programs were displaced by alignment and what types of variation these changes represented. We therefore projected expression through the row-normalized coupling and calculated an integration transport score (ITS) for each gene (Fig. 3a). The score measures the average standardized expression displacement induced by the fitted correspondence. ITS complements differential expression statistics by incorporating the distribution of matched cell populations encoded by the coupling.

**Fig. 3.**
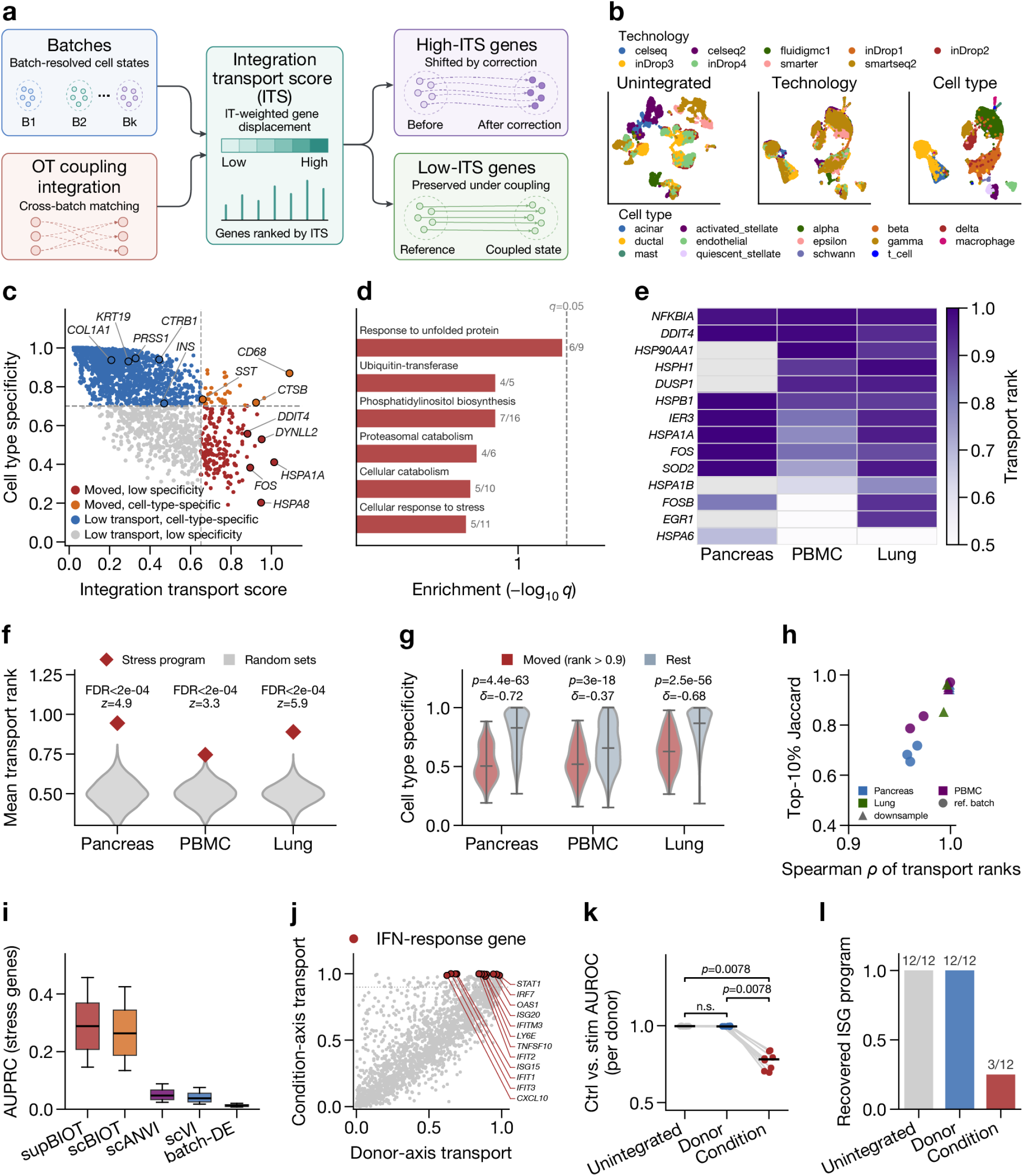
An explicit coupling makes batch integration auditable at gene resolution. **a**, Conceptual framework for the integration transport score, which ranks genes by coupling-weighted displacement and distinguishes genes shifted by correction from those preserved under coupling. **b**, Pancreas representations before and after integration. **c**, Integration transport score versus cell-type specificity. **d**, Gene Ontology enrichment among moved genes. **e**,**f**, Recurrence and random-set enrichment of a stress program across three datasets. **g**, Specificity of moved and other genes; two-sided Mann-Whitney tests with Cliff’s . **h**, Rank stability under reference changes and downsampling to 80% of cells. **i**, Comparative recovery of an externally defined tissue-dissociation and handling-stress signature on the nine-technology pancreas integration, comparing the transport score (supBIOT, scBIOT) with scVI and scANVI decoder attribution and per-batch differential expression. Boxes show the distribution of AUPRC values across 1,000 gene-level bootstrap iterations. **j**, Donor-axis and condition-axis transport in paired PBMC data. **k**, Per-donor control-versus-stimulation AUROC after no correction, donor correction or deliberately misspecified condition correction. Each point is one of eight donors, averaged over that donor’s cell types, with paired connectors and per-group medians. Brackets show paired two-sided Wilcoxon signed-rank tests across the eight donors (*n* = 8); the no-correction versus donor-correction comparison is marked n.s. because both sit at the AUROC ceiling and their difference is smaller than the donor-to-donor variation of the metric. **l**, Recovery of 12 prespecified interferon-stimulated genes (ISGs) after clustering.

In the pancreas benchmark, supBIOT mixed cells across nine technologies while preserving the major cell-type organization (Fig. 3b). To characterize the molecular changes induced by alignment, we jointly examined each gene’s transport score and cell-type specificity. This analysis distinguished genes that were minimally displaced, genes that were strongly displaced but broadly expressed, and genes that were strongly displaced despite high cell-type specificity (Fig. 3c). The latter category identifies potential alterations to cell-identity programs, whereas strongly displaced genes with low specificity are more consistent with shared technical or physiological variation.

Strongly displaced genes with low cell-type specificity included *HSPA1A, HSPA8, FOS, DYNLL2* and *DDIT4* (Fig. 3c). Functional enrichment showed that these genes were concentrated in the unfolded protein response, ubiquitin and proteasomal processes, cellular catabolism and antigen processing pathways (Fig. 3d). Thus, the inferred displacement reflected a coherent proteostasis and stress-response program rather than an arbitrary set of highly expressed genes.

This pattern was reproduced across pancreas, PBMC and lung integration analyses. Stress-response genes consistently received high transport ranks across datasets, although the contribution of individual genes varied by tissue (Fig. 3e). In each dataset, the mean transport rank of the stress program exceeded that of 5,000 matched random gene sets, with a false discovery rate (FDR) below 2 × 10^−4^ (Fig. 3f). Genes in the top transport decile also had significantly lower cell-type specificity than the remaining genes in all three datasets (Fig. 3g), supporting the preferential displacement of broadly shared programs rather than cell-type-specific molecular features.

Transport rankings were robust to reference selection and reduced sampling. Changing the reference batch or downsampling to 80% of cells produced high rank correlations and substantial overlap among genes in the top transport decile (Fig. 3h). These results indicate that ITS identifies reproducible molecular programs affected by alignment rather than variation driven primarily by reference choice or sampling noise.

To determine whether the displaced program reflected a known technical artifact rather than a pattern specific to these data, we used an externally defined tissue dissociation and handling stress signature^25^. This gene panel was fixed a priori and was not selected from the pancreas data. We used it as an independent positive set in the nine-technology integration and compared the coupling-intrinsic transport scores with scVI and scANVI decoder attribution^6,26^ and per-batch differential expression (Fig. 3i). Both transport variants recovered the dissociation program more accurately than the comparison methods in this dataset. The bootstrap median area under the precision-recall curve (AUPRC) was 0.29 for supBIOT and 0.25 for scBIOT, compared with 0.05 for scANVI, 0.03 for scVI and 0.01 for batch differential expression. The bootstrap distributions were wide because the dissociation-signature positive set was small, so the precision-recall estimates should be read alongside the more stable area under the receiver operating characteristic curve (AUROC) values. The corresponding point-estimate AUROC values were 0.96 for both transport variants, 0.87 for scANVI, 0.81 for scVI and 0.67 for batch differential expression. These results indicate that the program displaced by integration coincided with an independently defined technical stress signature, supporting the interpretation of ITS as an audit of nuisance-associated molecular displacement.

We next constructed a direct failure test using the Kang PBMC experiment, which crosses eight donors with control and interferon-stimulation^27^. When donor was used as the alignment axis, the condition contrast remained biological. When condition was deliberately specified as batch, the method was instead asked to remove the response of interest. Condition-axis transport correctly ranked the interferon program among the most strongly moved features (Fig. 3j). Deliberate condition correction lowered the per-donor control-versus-stimulation AUROC in all eight donors, from a median of 0.995 under donor correction to 0.781 (paired two-sided

Wilcoxon signed-rank *P* = 0.008, *n* = 8 donors, against either control; Fig. 3k). Donor correction, in contrast, left the AUROC at the ceiling (median 0.995 against the uncorrected 0.997), a difference smaller than the donor-to-donor variation of the metric and therefore not meaningful. A clustering and marker analysis recovered only 3 of 12 prespecified interferon genes after this misspecified correction, compared with 12 of 12 without correction (Fig. 3l). Because the underlying expression matrix was unchanged, this loss arose from the geometry used for clustering rather than from overwritten gene values.

Together with the dissociation-signature benchmark, this control illustrates how coupling-derived transport scores should be interpreted. When a strongly displaced program has low cell-type specificity, as observed for the proteostasis and dissociation-stress genes, its displacement is consistent with the alignment down-weighting shared technical variation. When a strongly displaced program is instead cell-type- or condition-specific and experimentally expected, as observed for the interferon response under misspecified correction, the same score indicates that the alignment design may be suppressing biology of interest. Coupling-derived transport scores therefore provide gene-level evidence for auditing an alignment and identifying cases in which its design should be reconsidered. This provides a second sense in which scBIOT is informed, complementing the label-guided transport used by supBIOT.

### A shared coupling resolves immune-state differences in HNSCC

We next asked whether the fitted coupling could localize disease-associated states after OT-based integration. We assembled a CD45-positive HNSCC atlas from GSE139324 and GSE164690^28,29^. Differences in sampling and cohort composition required integration to align shared immune states across the two studies. After quality control and integration, the tissue-restricted atlas contained 124,506 immune cells from 49 samples, including 5 healthy tonsils, 14 human papillomavirus (HPV)-positive tumors and 30 HPV-negative tumors (Extended Data Fig. 1a-d).

We calculated signed gene transport scores (GTS) and per-cell transport responses for three prespecified contrasts. Both healthy-to-tumor contrasts concentrated their largest responses in tumor-associated macrophage and monocyte states (Fig. 4a-c). By contrast, the comparison between HPV-negative and HPV-positive tumors localized strongly to the B-cell compartment. A two-axis summary separating the shared tumor effect from the HPV-associated effect placed germinal-center B (GC-B) cells and macrophages at opposing extremes (Fig. 4c,d). The coupling therefore nominated specific cellular compartments rather than returning only a list of cohort-level differentially expressed genes (Extended Data Fig. 1e,f).

**Fig. 4.**
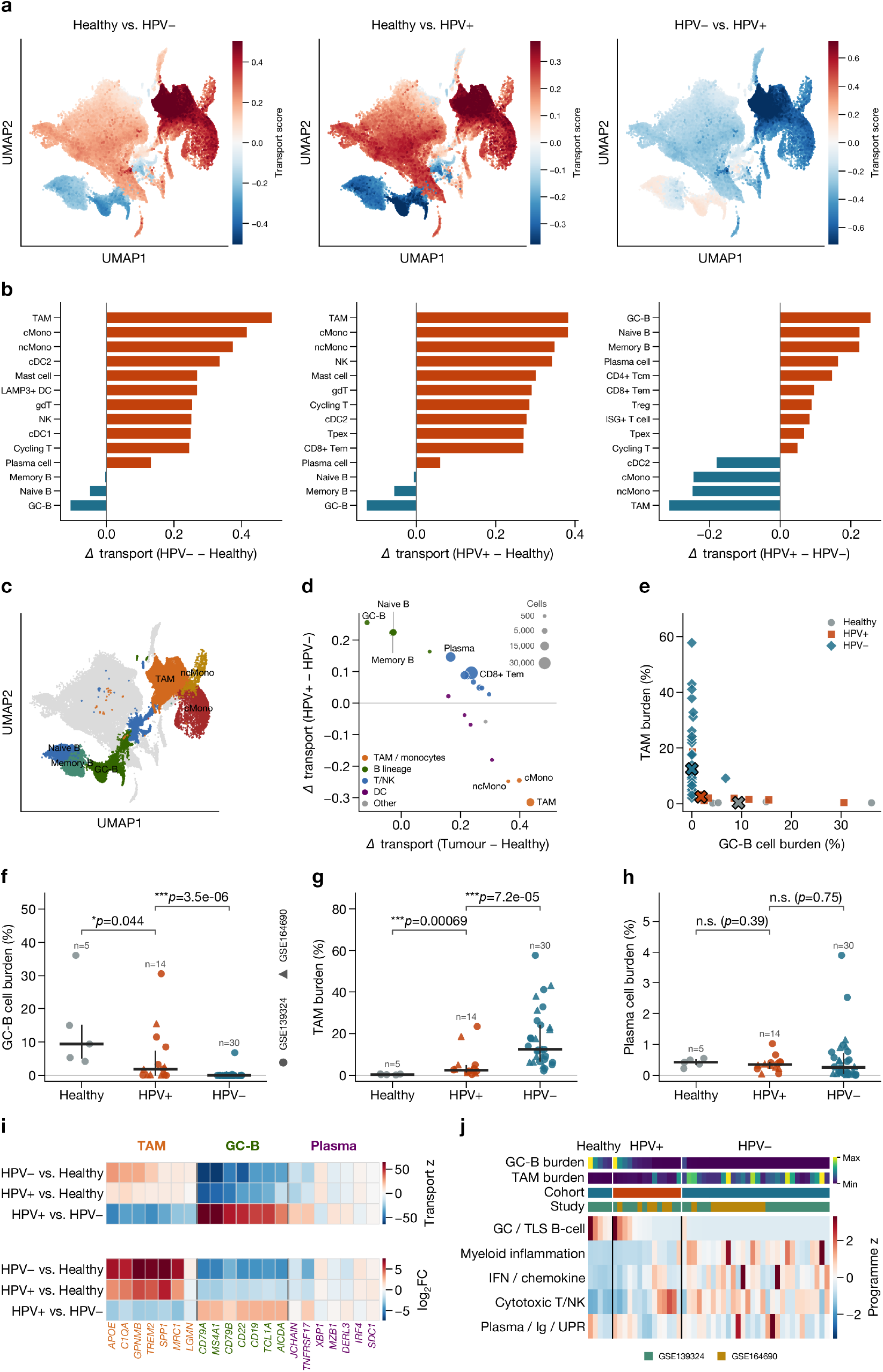
Transport scoring resolves reciprocal GC/TLS B-cell and macrophage states in HNSCC. **a**,**b**, Per-cell transport responses and cell-type impacts for three cohort contrasts. **c**,**d**, Integrated immune atlas and two-axis transport map. **e**, Per-sample GC-B and macrophage burdens. **f-h**, GC-B, macrophage and plasma-cell burdens across 5 healthy, 14 HPV-positive and 30 HPV-negative samples. Bars show median and interquartile range; brackets show two-sided Mann-Whitney tests. Marker shape denotes the source study (circle, GSE139324; triangle, GSE164690), shared across **f-h**; healthy samples derive from GSE139324 only. **i**, Gene-level signed transport and log_2_ fold-change matrices for macrophage (TAM), GC/TLS B-cell and plasma/Ig/UPR gene blocks, the same three populations quantified in **f-h**. TAM-block genes are restricted to those enriched in the annotated TAM compartment over both classical and non-classical monocytes. **j**, Program scores across samples. Cells localize signals but are not replicates. Absolute plasma-cell burden does not differ between tumor cohorts (*P* = 0.75).

To assess these cell-level patterns, we tested them statistically at the sample level. HPV-positive tumors contained a greater GC-B burden than HPV-negative tumors. The median abundance was 1.85% versus 0.00% of immune cells, with GC-B cells detected in only 8 of 30 HPV-negative tumors compared with 13 of 14 HPV-positive tumors (two-sided Mann-Whitney *P* = 3.5 × 10^−6^, Fig. 4e,f). Macrophage burden showed the reciprocal pattern and was greater in HPV-negative tumors, with a median abundance of 12.48% versus 2.51% of immune cells (*P* = 7.2 × 10^−5^, Fig. 4e,g). Absolute plasma-cell burden did not differ between the tumor cohorts, with median abundances of 0.35% and 0.26% of immune cells (*P* = 0.75, Fig. 4h). This result suggests that the HPV-associated B-cell signal was specific to the GC-B state rather than reflecting a general increase in plasma-cell differentiation. Sample-level program scores recapitulated the broader contrast (Fig. 4j). The directions of the GC-B and macrophage effects were reproduced within both source studies and exceeded those observed under sample-level label permutation (Extended Data Fig. 1g,h), reducing the likelihood that either result was driven by a single cohort or study. Gene-level transport and expression effects further distinguished a macrophage (TAM) program from a germinal-center/tertiary lymphoid structure (GC/TLS) B-cell program (Fig. 4i). The same genes separated along both healthy-versus-tumor axes, with TAM genes elevated and GC/TLS B-cell genes reduced in both contrasts (Extended Data Fig. 1i,j). To remove study-of-origin confounding, we repeated the healthy-versus-tumor comparison within GSE139324 alone, which provided all five healthy tonsils and 26 tumor samples. Myeloid-inflammation burden remained higher in tumor (median 13.2% versus 0.5% of immune cells, Cliff’s *δ* = 0.97, two-sided Mann-Whitney *P* = 4.7 × 10^−5^) and GC/TLS B-cell burden lower (median 0.0% versus 9.4%, *δ* = −0.83, *P* = 2.6 × 10^−3^). Because healthy tissue was available from only this study, external replication remains absent and we do not treat these healthy-versus-tumor effects as independently validated. The macrophage gene block was restricted to genes enriched in the annotated TAM compartment over both classical and non-classical monocytes, so that the gene-level evidence refers to the same population quantified at the sample level in Fig. 4g.

Together, these results show how a retained coupling can provide a common basis for atlas integration and disease-state inference. The same fitted correspondence that aligned the cohorts localized disease-associated differences to specific immune states, connected these states to implicated genes and programs and generated hypotheses for sample-level testing. scBIOT therefore does not replace conventional abundance or expression models. Instead, it makes their biological interpretation traceable to the cross-study correspondences used to construct the joint atlas.

### Prototype transport scales coupling analysis to 100 million cells

Cell-by-cell transport is infeasible for emerging perturbation atlases containing tens of millions of profiles. scBIOT therefore supports density-weighted prototype transport, in which cells are summarized into local states, OT is solved between prototypes and coupling-derived displacements are interpolated back to individual cells. We evaluated this strategy in the Tahoe-100M perturbation atlas^30^, analyzing 95.6 million cells after filtering across 50 cancer cell lines, 379 drugs and 26 mechanism-of-action groups (Extended Data Fig. 2).

Despite compression into prototypes, source-to-target occupancy remained structured. For instance, under ixazomib treatment, source states were preferentially routed to a limited set of target states rather than distributed diffusely across the atlas (Extended Data Fig. 2a).

Dominant target occupancy varied among mechanism groups when drugs were treated as replicates (Extended Data Fig. 2b), and program shifts recovered the observed differences between source and perturbed states (Extended Data Fig. 2c). These results show that prototype transport preserves interpretable state routing at a scale far beyond that accessible with an explicit cell-by-cell coupling.

We next compared the coupling-derived decomposition with marginal differential expression at this scale. For each perturbation-response program, we ranked genes by either marginal differential expression or coupling-specific GTS and assessed how well each ranking reconstructed the observed source-to-target routing (Extended Data Fig. 2d). Across programs, GTS-ranked genes reconstructed the routing at least as well as differentially expressed genes and more completely for several programs. Thus, the GTS ranking was not identical to the marginal differential-expression ranking. Because routing in this relatively homogeneous cell-line atlas was already largely reconstructable from differential expression, this analysis does not establish that transport scoring reveals biology beyond differential expression. Instead, it supports the narrower but important scalability claim that scBIOT can preserve and interrogate state-to-state routing across an atlas of 100 million cells through weighted prototype transport.

### Unified multiomic transport separates coordinated and modality-biased programs

We next tested whether a common transport geometry could distinguish biological programs with concordant cross-modal dynamics from those showing modality-specific divergence. We analyzed 10,862 erythroid-lineage cells with paired RNA expression and assay for transposase-accessible chromatin (ATAC) gene activity from bone marrow mononuclear cells (BMMCs)^31^. A separate cellular indexing of transcriptomes and epitopes by sequencing (CITE-seq)^32^ arm contributed 9,365 erythroid-lineage cells. Because the two arms contained different cells, they were linked through shared RNA, and protein-to-RNA comparisons were restricted to 16 mapped surface markers rather than same-cell correlations.

The joint representation aligned the paired multiome RNA and ATAC views with the RNA bridge from the separate CITE-seq arm while preserving the ordered progression from progenitors through erythroid maturation (Fig. 5a-c). Across the 16 surface markers, protein and RNA transport ranks were strongly concordant (Spearman *ρ* = 0.82, *n* = 16 markers, *P* = 1.1 × 10^−4^, Fig. 5d). This agreement indicated that the shared geometry recovered consistent marker-level dynamics across unpaired experimental arms. The limited marker set and absence of cell-level correspondence define the scope of this conclusion.

**Fig. 5.**
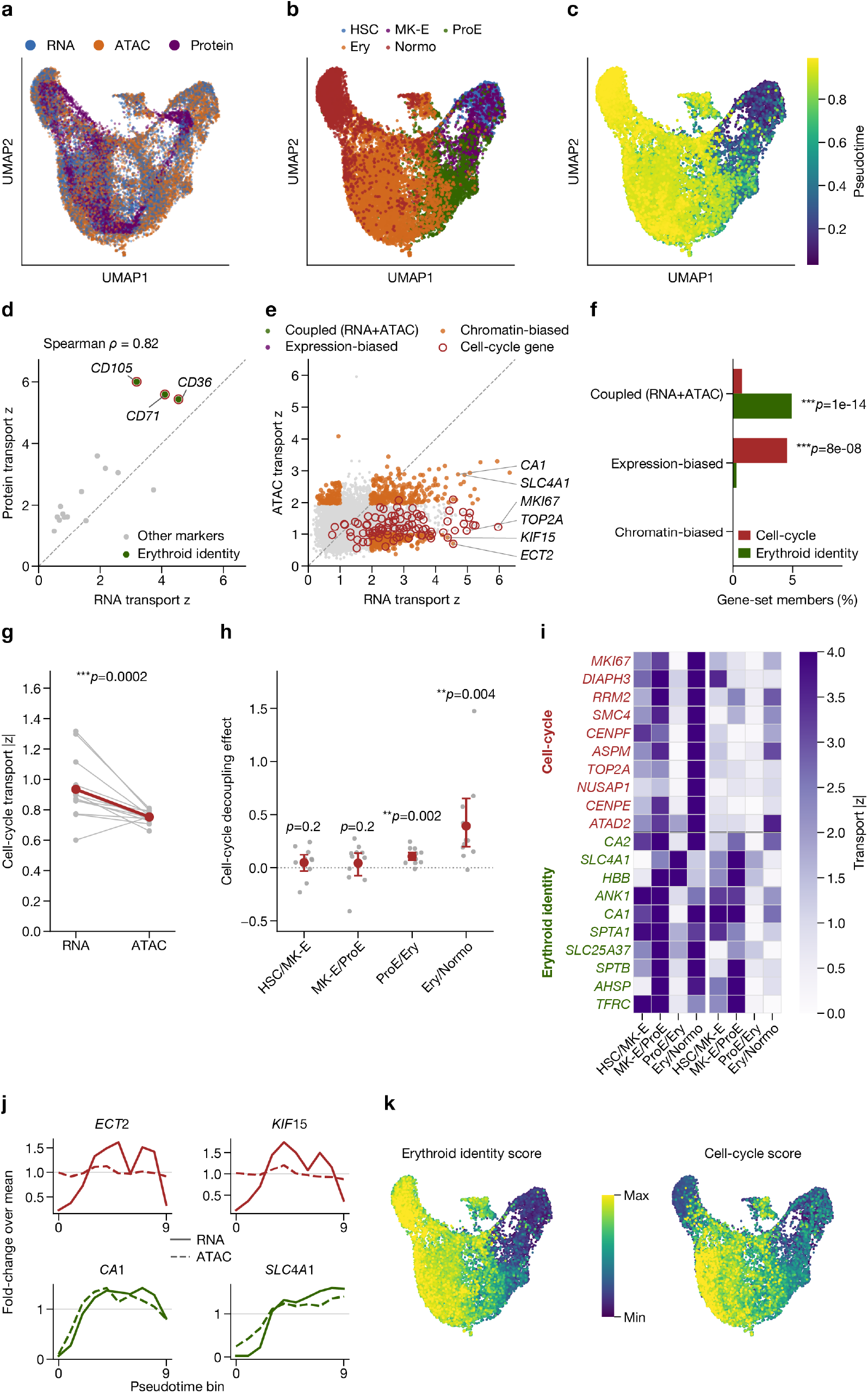
Multiomic transport separates lineage identity from expression-biased cell-cycle dynamics. **a-c**, Joint modality, erythroid-stage and pseudotime representations. **d**, Cross-arm protein and RNA transport for 16 mapped markers. Protein measurements come from different cells and are interpreted only at mapped-marker level. **e**, Paired-cell RNA and ATAC transport across 11,461 genes. **f**, Program enrichment within coupled and modality-biased classes. **g**, RNA and ATAC cell-cycle transport across 13 samples. The baseline-adjusted RNA-minus-ATAC effect was positive in all 13 samples (two-sided Wilcoxon signed-rank *P* = 2.4 × 10^−4^, *n* = 13). **h**, Baseline-adjusted effects by stage transition with bootstrap 95% confidence intervals and BH-adjusted tests. **i-k**, Transition heatmaps, gene trends and program maps.

The paired RNA and ATAC analysis revealed a different pattern. Across 11,461 genes, RNA and ATAC transport magnitudes were only weakly correlated (*ρ* = 0.19, Fig. 5e), indicating that concordant cell-state ordering did not imply uniform agreement across molecular programs. Genes with high transport in both modalities were strongly enriched for erythroid identity (hypergeometric *P* = 1.1 ×10^−14^), whereas genes with expression-biased transport were enriched for cell-cycle functions (hypergeometric *P* = 8.4×10^−8^, Fig. 5f). These are gene-level enrichment tests; the reproducible sample-level evidence for the cell-cycle asymmetry is given below. The coupling therefore separated a coordinated lineage program from a modality-biased dynamic program.

This distinction was reproducible across sequencing samples. The baseline-adjusted RNA-minus-ATAC cell-cycle transport effect was positive in all 13 sequencing samples (two-sided Wilcoxon signed-rank *P* = 2.4 × 10^−4^, *n* = 13), whereas raw cell-cycle transport was higher in RNA than in ATAC in 12 of 13 samples (Fig. 5g). Transition-specific estimates localized this difference primarily to proerythroblast-to-erythroblast and erythroblast-to-normoblast maturation (Fig. 5h), rather than to a uniform offset across the lineage. Gene-level heat maps and smoothed trends identified the genes contributing to these transitions, and program maps localized the effect along pseudotime (Fig. 5i–k).

A separately constructed B-lymphoid analysis reproduced expression-biased cell-cycle transport across 13 sequencing samples, but not the full erythroid lineage-identity pattern (Extended Data Fig. 3)^31^. This result is consistent with lineage programs being context specific, whereas the RNA and ATAC asymmetry of cell-cycle dynamics was shared across the two lineages. Together, these analyses show that a unified transport geometry can preserve a shared cell-state trajectory while distinguishing coordinated cross-modal programs from modality-biased programs. Although this framework does not establish whether one modality causally affects another, it identifies where modalities agree and where their contributions diverge without fitting separate correspondence models for individual programs.

### Spatiotemporal transport localizes developmental programs

Finally, we extended the retained-coupling strategy to tissue development using the Mouse Organogenesis Spatiotemporal Transcriptomic Atlas (MOSTA), a Stereo-seq atlas^33^. The analysis comprised 520,815 cells from eight mouse embryonic stages spanning E9.5 to E16.5. Expression was first represented in a stage-integrated latent space. Ordinal time and robustly standardized spatial coordinates were then introduced only during forward OT between adjacent stages. This design allowed developmental correspondence to incorporate spatial context without imposing spatial location on the initial expression integration (Fig. 6a).

**Fig. 6.**
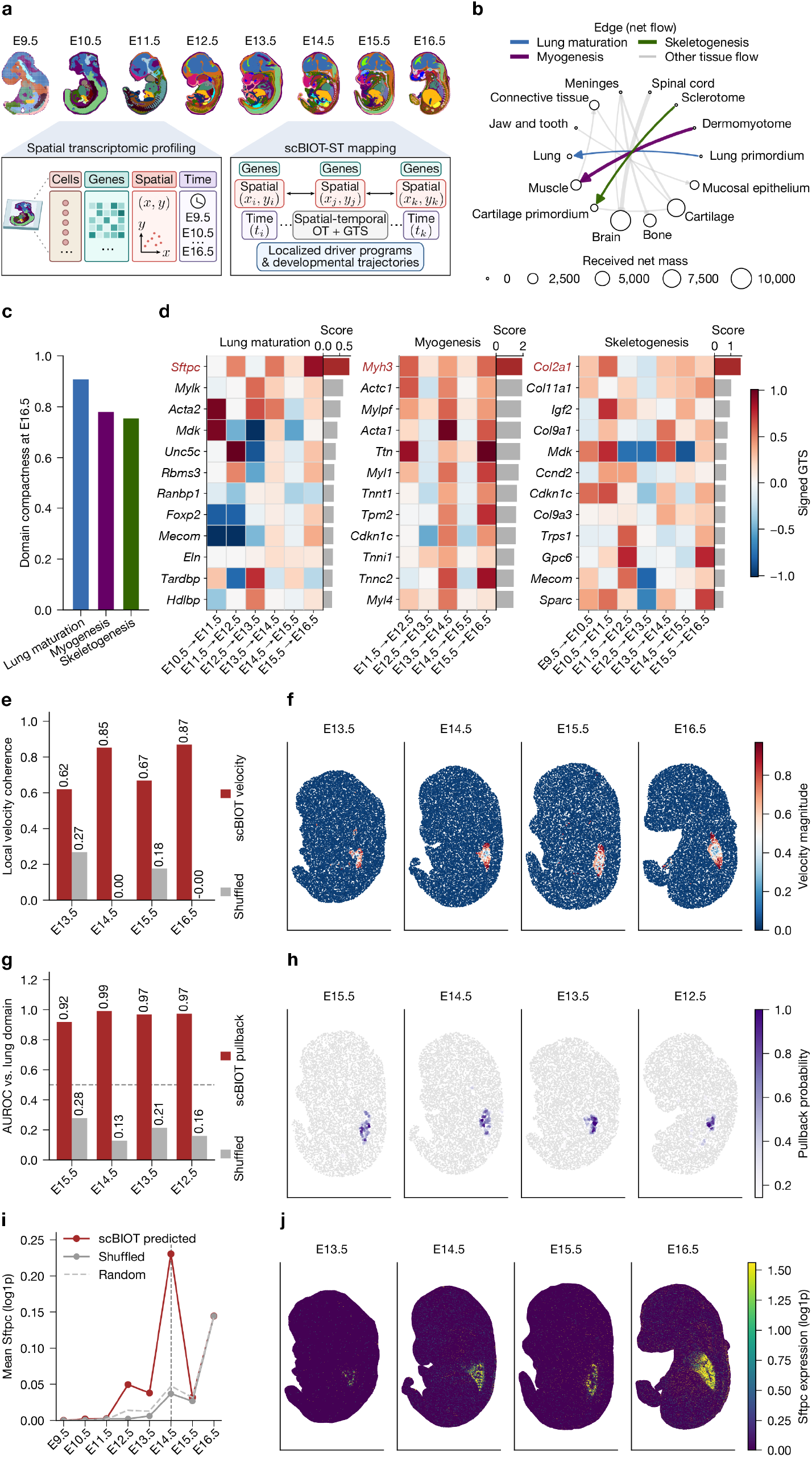
Spatiotemporal transport localizes developmental and gene-level programs. **a**, MOSTA analysis schematic. **b**, Tissue-domain net-flow network. **c**, Spatial compactness of three E16.5 target domains. **d**, Lineage-restricted signed GTS for lung, muscle and skeletal programs. **e**,**f**, Lung-region coherence and projected latent-velocity magnitude relative to shuffled controls. **g**,**h**, Lung-domain identification and spatial pullback from an E16.5 *Sftpc*-high target. **i**,**j**, *Sftpc* expression among transport-predicted cells and its direct spatial expression.

We compared scBIOT-ST with non-spatial scBIOT, moscot-spatial and moscot^4^, Waddington-OT^9^ and Sinkhorn-OT^34^ across four real and two simulated datasets. scBIOT-ST or non-spatial scBIOT achieved the highest or joint-highest label-transfer accuracy across the displayed benchmarks, the two scBIOT variants produced the lowest mean normalized entropies, and scBIOT-ST had the shortest recorded runtime (Extended Data Fig. 4a-c). These results illustrate the unified design of scBIOT, in which adjacent-stage couplings support domain transfer, gene prioritization, projected change and backward target localization within a single internally consistent framework.

The resulting domain-flow network recovered several anatomically coherent transitions, including Lung primordium-to-Lung, Dermomyotome-to-Muscle and Sclerotome-to-Cartilage primordium (Fig. 6b). Their E16.5 target domains were spatially compact (Fig. 6c), with domain-compactness scores of 0.91, 0.78 and 0.75 on a scale from 0 to 1 for the lung, muscle and skeletal targets, respectively. These values indicated that each flow terminated in a spatially coherent tissue region rather than being dispersed across the embryo. An additional analysis identified a neural-domain Spinal cord-to-Brain flow (Extended Data Fig. 4d,e). These couplings describe redistribution between annotated domains and do not directly observe migration or clonal fate.

We next asked whether the same couplings could identify molecular programs associated with each flow. Lineage-restricted signed GTS recovered *Sftpc* for lung development, *Myh3* and other sarcomere genes for muscle, and *Col2a1* together with cartilage collagens for the skeletal transition (Fig. 6d). Detection, false discovery rate and lineage-enrichment filters prevented broadly expressed genes from dominating the rankings. In the neural analysis, the same procedure highlighted *Tubb2b* (Extended Data Fig. 4e). Thus, domain-level mass redistribution and gene-level developmental programs were derived from the same adjacent-stage correspondence.

To test temporal composition more directly, we defined an Sftpc-high E16.5 lung target and composed row-stochastic couplings backward across adjacent stages. Around the developing lung, projected latent displacement was more coherent than in target-shuffled controls (Fig. 6e,f). The composed pullback assigned high probabilities to the lung domain at earlier stages. At E14.5, it identified lung cells with an AUROC of 0.99, compared with 0.13 after target-label shuffling (Fig. 6g,h). Sftpc expression was higher among cells assigned high transport-derived ancestry probabilities than among matched controls (Fig. 6i,j).

We applied the same adjacent-stage strategy to an axolotl telencephalon injury time course profiled by Stereo-seq^35^. The analysis recovered the reported injury-induced progenitor cascade, in which reactive ependymoglia give rise to regeneration intermediate progenitors and subsequently to neuroblasts. The recovered lineage was spatially confined to the injured hemisphere and temporally ordered across days after injury. Lineage-restricted GTS further nominated regeneration-associated genes, including *GLUL* (Extended Data Fig. 5). Together, these analyses indicate that the retained-coupling strategy is applicable across developmental and regenerative contexts. The same adjacent-stage formulation resolved both normal mouse organogenesis and axolotl brain regeneration while linking domain-level mass redistribution, lineage-restricted gene programs and backward temporal localization.

## Discussion

scBIOT establishes an informed, scalable and unified framework for single-cell atlas integration by making the fitted correspondence an explicit and inspectable analytical object. Its contribution extends beyond generating corrected coordinates. The supervised formulation incorporates known cell identities into the transport geometry to discourage biologically implausible matches. Across seven heterogeneous transcriptomic and spatial benchmarks, evaluated on identical cells and labels for every method, supBIOT achieved the highest macro and weighted label-transfer F1 in each one while preserving biological conservation. This balance is important because globally well-mixed embeddings can still yield unreliable annotations, particularly for rare populations. After alignment, the integration transport score reveals which molecular features were displaced by the fitted correspondence, providing a gene-level audit not available from embedding-level summaries alone. Its recovery of a reproducible dissociation-associated stress program across pancreas, PBMC and lung datasets showed that the score can identify shared technical signals. Density-weighted prototypes further preserved interpretable state-level transport across 95.6 million cells, extending coupling-based analysis to atlas scale. In multiomic analyses, the coupling separated programs with concordant cross-modal dynamics from those biased toward a single modality. In spatiotemporal analysis, scBIOT variants matched or exceeded the comparison methods on label-transfer accuracy while localizing developmental transitions and prioritizing candidate developmental genes. Together, these results connect the central design principles of scBIOT to measured performance and capabilities in reference mapping, label transfer, gene-level auditing, atlas-scale analysis, multiomic comparison and spatiotemporal inference.

Previous single-cell OT methods established transport-based inference for temporal and lineage analysis^4,9,10^, spatial analysis^13,15,16^, multimodal alignment^14,18^ and perturbation modeling^19,20^. The central methodological contribution of scBIOT is an atlas-integration framework in which explicit, analysis-specific OT couplings support both alignment and subsequent inference. The coupling used for mapping is retained to derive mapped positions, to define the aligned embedding in which label probabilities are estimated, to quantify gene-level displacement, to characterize state-level correspondences and to support downstream biological contrasts. By combining these functions with biologically informed transport and density-weighted prototypes, scBIOT connects mapping, molecular auditing and atlas-scale downstream analysis within a common computational framework. This correspondence-centered design directly links biological findings to the alignment that generated them and preserves information that would otherwise be lost when integration and downstream inference use separate representations. scBIOT therefore treats atlas integration as an explicit and inspectable coupling rather than an opaque preprocessing step.

A second methodological advance is density-weighted prototype transport, which makes coupling-based inference tractable without relinquishing state-to-state correspondence. Unlike conventional subsampling, the prototypes represent local cellular states and retain their empirical masses, allowing transport normalization, state routing and gene-level scoring to remain consistent with the cell-level formulation. In the Tahoe-100M atlas, this strategy preserved interpretable routing across 95.6 million cells, 50 cancer cell lines and 379 perturbations. Structured prototype occupancy revealed how perturbations redistributed cells among target states, while transport-derived programs linked these redistributions to specific molecular features. These scores complemented rather than duplicated marginal differential expression. Differential expression better captured overall perturbation magnitude, whereas gene transport scores identified features associated with state redistribution. Prototype transport thus extends explicit correspondence analysis to atlas scale while preserving a distinct interpretation of the transported mass.

A third advance is the retention of coupling-derived correspondences as a common basis for inference across conditions and modalities. In HNSCC, the correspondences used to integrate cohorts also linked tumor-associated responses to macrophage states and HPV-associated responses to GC-B states, while ranking the gene programs underlying these contrasts. These hypotheses were supported by sample-level analyses of cell abundance and gene programs. In the multiomic analysis, a shared transport geometry recovered concordant protein and RNA dynamics while distinguishing coordinated erythroid programs from cell-cycle dynamics that were stronger in RNA than in chromatin. This RNA and ATAC asymmetry was reproduced across sequencing samples and in an independently constructed B-lymphoid analysis. These applications demonstrate a specific advantage of the unified formulation. Cell-state correspondences and gene-level interpretations remain internally consistent because both are conditioned on the same transport geometry rather than reconstructed using separate models for each modality or biological program.

Spatiotemporal analysis further tested this correspondence-centered design by extending it from single-step to composable couplings. scBIOT-ST introduces spatial and ordinal temporal information into forward transport between adjacent developmental stages and then composes the resulting row-stochastic couplings across time. The same transport paths consequently support tissue-domain transfer, projected change, backward target localization and prioritization of candidate developmental genes. Adjacent-stage couplings recovered anatomically coherent lung, muscle and skeletal transitions together with lineage-associated genes including *Sftpc, Myh3* and *Col2a1*. Composed couplings traced an *Sftpc*-high lung target backward through development and localized its earlier-stage lung domain more accurately than target-label-shuffled controls. These results highlight composable transport as a defining advance of scBIOT-ST. By embedding spatial and ordinal temporal structure into stage-to-stage couplings and composing them across development, a single correspondence model both localizes where transitions occur and traces their targets backward through time, providing biological information not available from single-step alignment.

These advances also delineate the limits of the framework. OT solutions remain sensitive to the representation, transport cost, regularization and marginal penalties, and concentrated transport does not by itself establish biological validity. Supervised transport depends on the quality and completeness of reference annotations, although unbalanced transport allows states without a close reference counterpart to retain unmatched mass. Prototype transport can reduce noise in the empirical data distribution but does not exactly reproduce the full cell-level coupling. Finally, transport-derived relationships remain associative and do not establish causality.

Taken together, scBIOT advances single-cell atlas integration by treating the fitted OT coupling as the central analytical object that links alignment to subsequent inference. The framework combines biologically informed transport that constrains implausible correspondences, integration transport scores that audit molecular displacement induced by alignment, density-weighted prototype transport that preserves interpretable mass routing at atlas scale and composable spatiotemporal couplings that propagate correspondences across developmental stages. Because mapped coordinates, label probabilities, state-level relationships, gene-level interpretations and downstream biological contrasts are derived from the same transport geometry, each conclusion remains directly traceable to the alignment that generated it. scBIOT therefore extends atlas integration beyond coordinate correction into an explicit, scalable and testable model of cellular correspondence that supports reference mapping, molecular auditing, multimodal comparison and spatiotemporal inference within a common computational framework.

## Methods

### Computational overview

Let 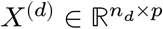 denote the mean-centered, normalized feature matrix for domain *d*, where a domain can be a batch, study, condition, modality or developmental stage. scBIOT first learns a shared *k*-dimensional geometry with a tied linear autoencoder whose encoder and decoder are

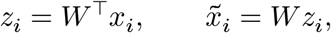

with loadings *W* estimated by minimizing

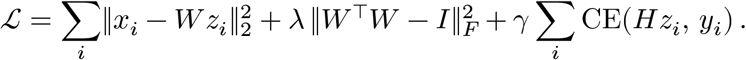

Here, *I* is the *k* × *k* identity matrix, *λ* weights the orthogonality penalty and CE denotes class-weighted cross-entropy loss. The reconstruction and orthogonality terms make *W* behave like principal component analysis (PCA) loadings. The final cross-entropy term is active only in supervised (label-transfer) fits, where a linear head *H* classifies the reference label *y*_*i*_ and shapes the shared geometry without breaking its linearity. Unlabeled cells are ignored by this term, and *γ* = 0 in unsupervised fits. The head bias is omitted from the notation.

The tied weights keep the representation directly connected to the original feature space, and the orthogonality penalty discourages redundant latent axes. The implementation operates on sparse normalized matrices, and batch effects are handled downstream by the OT integration step. All random seeds and figure-specific preprocessing choices are recorded in the analysis scripts.

For source representation *Z*^(*s*)^ and target or reference representation *Z*^(*t*)^, the default ground cost is squared Euclidean distance, 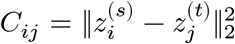. With normalized source and target weights *a* and *b*, scBIOT estimates an entropically regularized unbalanced coupling^34,36^

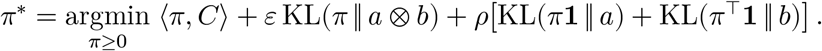

Here, **1** is a vector of ones, *a* ⊗ *b* is the independent product measure and KL denotes Kullback-Leibler divergence. The parameter *ε* controls coupling smoothness, and *ρ* controls deviation from the specified marginals. The unbalanced penalties allow unmatched or depleted states while still discouraging arbitrary mass loss. Costs were scaled within each analysis before optimization so that regularization had a comparable interpretation across domain pairs.

For mapping, the coupling is row normalized as 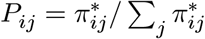, and the barycentric projection of source cell *i* into the target geometry is

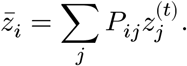

Rather than adopting 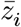 directly, scBIOT integrates by moving each cell toward its coupled target while preserving local neighborhood structure, minimizing

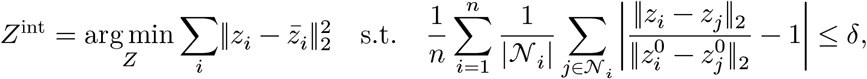

where *N*_*i*_ denotes the set of *k*-nearest neighbors of cell *i* in its original embedding *z*^0^ (distances stabilized with a small constant) and *δ* bounds the mean admissible relative distortion; a matching upper-quantile cap prevents individual cells from tearing. The alignment term removes batch geometry, while the distortion budget keeps transport from collapsing or tearing biological structure. The coupling supplies coordinates. Specifically, *P* defines the barycentric targets 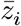 that drive the integration step above, such that each cell’s position in the shared aligned embedding is determined by transport. The coupling does not carry label mass, and labels are not transported along *π*^∗^. Instead, labels are assigned in a separate readout step after alignment. A multinomial logistic regression classifier is fitted to the labeled reference cells in the shared aligned embedding and then applied to the query cells. The class with the highest predicted probability is assigned as the transferred label, and the corresponding probability is reported as the confidence.

scBIOT operates in an unsupervised manner by default. When reference annotations are available, the supervised variant (supBIOT) uses them to keep distinct cell identities from being merged during alignment and to carry labels onto the query. Identity enters at two stages. The shared encoder is trained with the class-supervised head above, and the structure-preserving integration adds a label-guided displacement

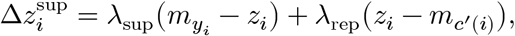

pulling each labeled cell toward its reference-class centroid 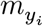 and away from the nearest competing centroid 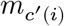. The two coefficients are not independent parameters. Both are tied to a single supervision level *s* ∈ [0, 1] by *λ*_sup_ = 0.8 *s* and *λ*_rep_ = 0.25 *s*, so *s* scales the displacement as a whole while the fixed 0.8 : 0.25 ratio keeps centroid attraction dominant over repulsion; *s* = 0 recovers the unsupervised behavior. We did not tune *s* per dataset. All supervised analyses reported here use the package default *s* = 0.5, and the same value is used for every benchmark, so reported label-transfer accuracies reflect a single fixed setting rather than a per-dataset search.

Query cells carry no reference label, so *λ*_sup_ = *λ*_rep_ = 0 for them and 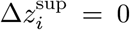; their coordinates are set entirely by expression geometry and the coupling-driven alignment. Their labels are then obtained from the post-alignment classifier readout described above, so no query label is imposed by the displacement term. This preserves known populations and separates confusable states without requiring the source and reference label sets to match.

For datasets too large to store a cell-by-cell cost matrix, scBIOT replaces cells with prototypes computed independently within each batch by weighted mini-batch *k*-means (up to 2,048 prototypes per batch by default). Each cell is weighted by its local sparsity, the mean Euclidean distance to its 15 nearest neighbors normalized by the global mean, so that sparse and bridging regions are not under-represented. Prototype mass equals the number or summed sampling weight of its assigned cells. OT is solved between prototypes, and each cell’s displacement is the inverse-distance-weighted average of the barycentric displacements of its 8 nearest prototypes. This approximation changes the resolution of the coupling, not its normalization or downstream definitions.

### Reference mapping benchmark

supBIOT (scbiot v1.2.1) was compared with Symphony (symphonypy v0.2.4 with harmonypy v0.0.10), scANVI (scvi-tools v1.4.1), scPoli (scArches v0.6.1 with scipy v1.13.1), Seurat (v5.5.1 with SeuratObject v5.4.0 and zellkonverter v1.22.0 under R v4.6.1) and an unintegrated PCA embedding as a baseline^5–8^. Because these packages pin mutually incompatible dependencies, each method was run in its own environment; harmonypy and scipy were pinned to the versions above because later releases removed interfaces that symphonypy and scArches require. Tasks included pancreas, PBMC, Tabula Muris Senis, Tabula Sapiens and Xenium colorectal, liver and ovarian cancer datasets^1,21–24^. Preprocessing was performed once per dataset and written to a frozen input that every method then read, so cells, features, reference/query split, labels and random seed were identical across methods by construction. Feature selection followed one rule fixed by assay type rather than by dataset: targeted panels were used in full, and whole-transcriptome datasets were reduced to the top 3,000 highly variable genes (seurat_v3 flavor on counts, computed within batch). The unintegrated baseline used 50 principal components for pancreas and PBMC and 100 for the remaining datasets. Xenium queries were additionally filtered as in the source protocol, retaining cells with at least 20 counts and below the 98th percentile of total counts and genes detected in at least 100 cells.

Integration performance was evaluated using scib-metrics v0.5.9. Biological conservation was assessed using isolated-label, silhouette, Leiden normalized mutual information (NMI), Leiden adjusted Rand index (ARI) and cell-type local inverse Simpson’s index (cLISI) metrics. Batch correction was assessed using batch-removal-adapted silhouette (BRAS), integration local inverse Simpson’s index (iLISI), k-nearest-neighbor batch-effect test (kBET), graph connectivity and principal component regression (PCR) comparison metrics. Metric values were not min-max scaled across methods. They were averaged within each group, and the displayed overall integration score was 0.6 times the biological-conservation mean plus 0.4 times the batch-correction mean. Each method produced query labels with its own native transfer procedure, so that every method is evaluated as intended by its authors. The unintegrated baseline used a multinomial logistic-regression classifier (scikit-learn) trained on labeled reference cells in the PCA embedding. Predictions were compared with the withheld query annotations using a single scoring implementation shared by all methods. A cell entered the evaluation if it belonged to the query split and carried a ground-truth label, a definition that depends only on the data; a method returning no prediction for such a cell was scored as incorrect rather than having the cell excluded, so every method was scored on exactly the same cells. Label-transfer performance was summarized by weighted and macro F1. Weighted F1 weights each class by its query-cell abundance. Macro F1 is the unweighted mean of class-specific F1 values. For each method, “Overall” in Fig. 2a–c is the unweighted arithmetic mean of the seven dataset-specific scores.

### Transport scores

Genes were filtered by detection, counts were normalized and log transformed, and highly variable genes were selected separately for each analysis. scBIOT derives two complementary gene-level scores from the fitted coupling. The integration transport score (ITS) measures the extent to which integration shifts a gene, whereas the gene transport score (GTS) attributes a source-to-target contrast to individual genes.

For each non-reference batch, log-normalized reference expression was projected through the row-stochastic coupling, 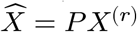, after standardizing each gene so that abundant genes did not dominate through scale alone. The ITS of gene *g* is the mean absolute projected shift,

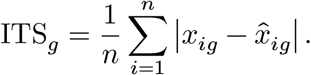

The score is non-negative; genes with larger alignment-associated differences have greater displacement, whereas conserved genes remain more stable. With several non-reference batches, ITS_*g*_ was computed per batch pair, averaged with equal weight across the valid batch pairs and converted to a within-set percentile rank.

The GTS instead uses the raw coupling *π*^∗^ between a source group and a target group and weights every matched pair (*i, j*) by its transported mass. The unsigned transport magnitude is

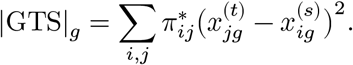

For signed analyses, the corresponding first-moment score, 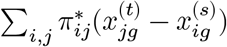, was computed in both transport directions. The reverse score was sign-aligned to the forward direction and the two null-standardized scores were averaged, 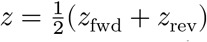. Scores were normalized by the pooled gene-specific variance and converted to *z* scores using a matched null generated by permuting target-cell indices (expression profiles) relative to the fixed coupling weights *π*^∗^, or using its analytic normal approximation. Writing 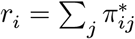 and 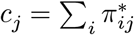 for the source- and target-cell marginals, the analytic form treats the signed score as a weighted sum of independent per-cell values, 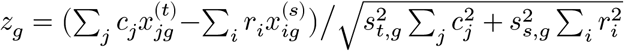, with 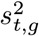 and 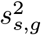 the per-gene target and source expression variances and two-sided 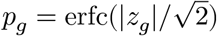. For ordered contrasts, per-step *z* scores were combined by Stouffer’s method, 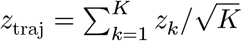 across the *K* adjacent-stage steps, which defines the reported *p* and FDR, whereas genes were ranked by the step magnitude 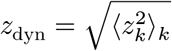.

Cell-type specificity was quantified using the *τ* index calculated from cell-type mean expression. Moved genes were defined as those with a transport rank above 0.9. Their specificity was compared with that of all other genes using a two-sided Mann-Whitney test and Cliff’s *δ*.

Gene Ontology (GO) over-representation used a hypergeometric test against the analyzed gene background. Benjamini-Hochberg (BH) correction controlled FDR. A declared stress program was tested in each dataset by comparing its mean transport-rank percentile with a null distribution of 5,000 random gene sets of the same size, each drawn without replacement from that dataset’s ranked genes, so the null was matched on set size alone. A one-sided empirical P value was computed as (1 + *b*)/(5,000 + 1), where *b* is the number of null sets whose mean rank equaled or exceeded the observed value, and BH correction was applied across the three datasets (pancreas, PBMC and lung). Transport-rank stability was assessed after reference-batch changes and downsampling to 80% of cells. We reported Spearman correlation and top-decile Jaccard overlap against the primary ranking.

For the Kang control^27^, donor transport was matched within condition, and condition transport was matched within cell type, preventing donor composition from defining the stimulation score. A deliberately misspecified fit then used condition as the batch variable. This control was specified to test whether the audit identified a known biological program when the integration design incorrectly requested its removal.

Control and stimulation were distinguished using logistic regression on the embedding coordinates, with out-of-fold AUROC from stratified fivefold cross-validation. To base the readout on independent biological replicates, the AUROC was computed within each donor and cell type and then averaged across cell types to give one value per donor, so that cell-type composition rather than the stimulation response was held out. The eight donor-level AUROC values were compared across the three integration settings with paired two-sided Wilcoxon signed-rank tests. The uncorrected and donor-corrected embeddings both retained the stimulation signal at the AUROC ceiling, where their median difference of about 0.001 was smaller than the donor-to-donor variation of the metric; that comparison was therefore reported as not significant rather than assigned a p-value. Marker recovery was evaluated using Leiden clustering followed by identification of the top 25 Wilcoxon markers from the unmodified expression values. The external dissociation signature was obtained from van den Brink et al.^25^. Method differences in AUPRC were assessed with a gene-level bootstrap. The full set of analyzed genes was resampled with replacement to its original size for 1,000 iterations, with the dissociation signature as the positive class and no stratification of positive and negative genes. The same 1,000 resampled index sets were applied to every method, giving a paired comparison, and AUPRC was recomputed for each method on each resample. Iterations whose resample contained no positive genes or only positive genes were discarded. Figure 3i shows the resulting per-method AUPRC distributions, and the median across iterations is reported in the text.

### HNSCC analysis

Raw HNSCC data were obtained from GSE139324 and GSE164690^28,29^. The integrated CD45-positive atlas comprised 198,702 cells and 22,164 genes. Tissue analyses were restricted a priori to 124,506 tonsil or tumor-infiltrating cells from 49 samples. Cells were retained if they had between 200 and 6,000 detected genes, a mitochondrial fraction below 20%, a ribosomal fraction below 50% and a complexity score (log_10_ genes per UMI, the ratio of log_10_ detected genes to log_10_ total counts) between 0.8 and 1.0. Doublets were predicted separately within each sample using `scanpy.pp.scrublet` (Scanpy v1.12.1, wrapping Scrublet v0.2.3) and excluded. Cell types were annotated by Leiden clustering of the integrated embedding, assigning each cluster from its lineage markers (for example germinal-center B cells from *RGS13, AICDA, MS4A1* and *MKI67* ; tumor-associated macrophages from *CD68, C1QC, APOE* and *CD14*; plasma cells from *MZB1, XBP1, JCHAIN* and *DERL3*), and grouped into B, myeloid, T/NK and other immune lineages. Disease analyses included three contrasts comprising healthy versus HPV-negative, healthy versus HPV-positive and HPV-negative versus HPV-positive samples. Gene transport scores were computed for 2,000 highly variable genes using signed transport aggregation and a permutation-based null distribution.

Per-cell disease scores were used to localize transport across the atlas, and cell-type impact was defined as the between-cohort difference in the mean score. The shared tumor axis was calculated by averaging the impacts of both tumor-versus-healthy comparisons, whereas the HPV axis contrasted HPV-positive and HPV-negative tumors. Samples were treated as the unit of cohort-level inference. Program activity was summarized per sample in two ways. Count-based burdens expressed each cell-type-defined program as a fraction of the sample’s total immune cells, namely the germinal-center/TLS B-cell and plasma-cell fractions, the combined tumor-associated-macrophage and monocyte fraction for myeloid inflammation, and the combined CD8+ effector-memory, NK and *γ δ* T-cell fraction for cytotoxicity. Expression-based programs (interferon/chemokine and plasma/immunoglobulin/UPR) were the per-cell mean of their marker panels averaged to a per-sample pseudobulk and z-scored across samples. Marker panels were *ISG15, MX1, IFIT1, IFIT3, STAT1, CXCL9, CXCL10* and *CXCL11* (interferon/chemokine); *PRDM1, IRF4, XBP1, MZB1, JCHAIN* and *SDC1* (plasma/immunoglobulin/UPR); *BCL6, AICDA, RGS13, MEF2B, CXCR4* and *CD83* (GC/TLS B-cell); *CD68, C1QA, C1QC, APOE, TREM2, SPP1, IL1B, CXCL8, S100A8* and *FCN1* (myeloid inflammation); and *GZMB, GZMK, NKG7, GNLY, PRF1, KLRD1* and *CD8A* (cytotoxic T/NK). Per-cell program scores used `scanpy.tl.score_genes`, the mean panel expression minus an expression-matched control gene set. All 49 tissue samples were retained, including samples with zero counts. Primary comparisons used two-sided Mann-Whitney tests. Robustness was assessed using study-stratified analyses and a sample-level permutation test that shuffled cohort labels across the tumor samples, restricted to the HPV-positive and HPV-negative groups (5,000 permutations), with a two-sided empirical P value from the fraction of permutations whose absolute median cohort difference matched or exceeded the observed value. To remove study-of-origin confounding from the healthy-versus-tumor comparison, program burdens were compared between the five healthy tonsils and 26 tumor samples of GSE139324 alone using two-sided Mann-Whitney tests and Cliff’s *δ* effect sizes.

### Tahoe-100M prototype-transport analysis

The Tahoe-100M analysis included 95.6 million cells retained after filtering across 50 cell lines, 379 drugs and 26 mechanism-of-action groups^30^. Plates were first integrated by centroid OT (up to 2,048 prototypes per plate). Within each cell line, Leiden clustering of the integrated embedding defined discrete states; vehicle-treated states served as sources and drug-treated states as targets. Entropic OT (Sinkhorn regularization 0.05) between their mean coordinates, with marginals set to the state cell fractions, yielded a row-normalized source-to-target occupancy matrix. Dominant target occupancy was defined as the largest target-state probability for each source state.

Program shift was calculated as the target-minus-source change after weighting target states by their coupling occupancy. Routing reconstruction (Extended Data Fig. 2d) quantified how well the observed per-program target shift was predicted from the top-ranked genes. For each response program, a ridge regression on the top marginal differential-expression or coupling-specific GTS genes was scored by drug-grouped five-fold cross-validated *R*^2^. Drugs, rather than drug-dose-cell-line combinations or individual cells, served as the replication unit for mechanism-group summaries.

### Multiomic analysis

Processed GSE194122 resources came from the NeurIPS multimodal integration benchmark^31^ and comprised a paired 10x Multiome arm and a separate CITE-seq arm. The erythroid Multiome subset contained 10,862 cells with same-cell RNA expression and ATAC gene activity over 11,461 shared genes. The independently constructed B-lymphoid replication contained 10,713 paired Multiome cells.

The CITE-seq erythroid arm contained 9,365 cells and 134 surface proteins. These cells were not paired to cells in the Multiome arm. CITE-seq RNA was restricted to the 10,899 genes shared with the erythroid Multiome data and provided the bridge between assay arms. Sixteen proteins had unambiguous mapped gene symbols and entered the protein-RNA comparison. Protein comparisons were therefore cross-arm marker analyses and were not interpreted as cell-level molecular correlations.

Multiome RNA expression, ATAC gene activity and CITE-seq RNA expression were normalized to 10,000 counts per cell and log transformed. Surface-protein abundance was centered log-ratio (CLR) transformed. Multiome RNA and CITE-seq RNA formed the reference expression view, and Multiome ATAC gene activity was projected into that space by the reference-fitted encoder. Treating assay view as the batch, scBIOT aligned the three views in a common latent geometry. The paired RNA and ATAC coordinates of each Multiome cell were averaged into one shared-state vector,

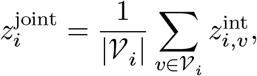

where *V*_*i*_ indexes the paired RNA and ATAC views of Multiome cell *i*. Each CITE-seq cell retained the coordinate of its own RNA view. No cell-level correspondence between the CITE-seq and Multiome arms was used.

Pseudotime was divided into ten quantile bins within each assay arm. On the paired Multiome geometry, GTS was evaluated separately for RNA expression and ATAC gene activity, giving permodality magnitudes 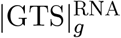 and 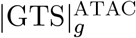. Protein transport was evaluated independently in the CITE-seq cells within the shared geometry. Dynamic transport magnitude was defined as the root mean square of the step-specific signed transport *z* scores. Concordance between protein and RNA was assessed across the 16 mapped markers using Spearman correlation. Because the scores were derived from different cells and assay arms, this analysis measured agreement in marker rankings rather than same-cell covariation.

Genes above the pooled 90th percentile were considered transported, and genes below the 35th percentile were considered untransported. Coupled, expression-biased and chromatin-biased classes followed from these declared cutoffs. Program enrichment used hypergeometric tests against the analyzed gene universe. A threshold-free Mann-Whitney test compared RNA-to-ATAC transport ratios for cell-cycle genes against other genes. Sequencing sample was the replication unit for whole-lineage and transition analyses. Baseline-adjusted RNA-minus-ATAC effects were tested with two-sided Wilcoxon signed-rank tests. Transition tests were BH corrected, and a sample bootstrap provided 95% confidence intervals.

### Spatiotemporal analysis

The MOSTA object contained 520,815 cells, 23,761 genes, eight stages and 45 tissue annotations^33^. Counts were normalized to 10,000 and log transformed. Two thousand highly variable genes entered embedding and transport scoring. A tied linear autoencoder produced the L2-normalized expression representation. Stages were aligned with centroid unbalanced OT, which did not receive spatial coordinates or time. Spatial coordinates were centered and robustly scaled within each stage; in the forward-transport cost their weight was scale-matched to twice the embedding’s mean per-dimension standard deviation, so the two spatial dimensions did not dominate the embedding dimensions. Each stage was summarized by 1,500 density-weighted centroids, adjacent stages were coupled by entropic unbalanced OT (regularization 0.05), and couplings were computed only between consecutive stages in increasing time.

The velocity of source centroid *c* was its barycentric displacement into the next stage,

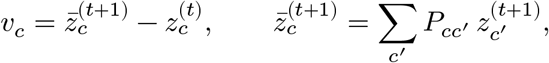

interpolated to cells by inverse-distance weighting over the 8 nearest centroids. Redistribution between annotated domains was summarized as the transported mass 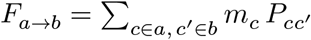 (*m*_*c*_ the source-centroid mass), with the net-flow network given by *F*_*a*→*b*_ −*F*_*b*→*a*_. Target-domain compactness at E16.5 was defined as 1 − ‖sd(**s**)‖_2_/*D*_bbox_, the L2 norm of the target-tissue cells’ spatial-coordinate standard deviation normalized by the embryo’s bounding-box diagonal (range 0 to 1, higher indicating a more spatially compact domain). Velocity coherence was the mean cosine similarity between each cell’s projected velocity and the mean velocity of its 15 nearest spatial neighbors, with a within-stage coordinate shuffle as the null. Projection into physical coordinates visualized latent developmental change and was not interpreted as physical migration.

Signed GTS used matched 6,000-cell subsamples for adjacent stages and an analytic Wald test. Genes required detection in at least 3% of cells. BH-adjusted *q* < 0.05 defined significance. Lineage-specific scores further required detection in at least 5% of lineage cells and expression enrichment of at least 0.20 log units. Genes were ranked by lineage enrichment after FDR filtering.

For lung pullback, an E16.5 *Sftpc*-high target was defined as the top 30% of E16.5 lung cells by *Sftpc* expression. Row-stochastic adjacent-stage couplings were composed across time, 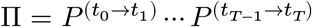, so that Π applied to the target indicator returns each early cell’s pullback (ancestry) probability; *Sftpc* expression was averaged over the top-decile ancestry cells at each stage. Controls included shuffled targets, temporal-only coupling, expression-matched random genes and lung-domain prediction at E14.5. These analyses provide coupling-derived ancestry probabilities rather than validated fate assignments. The benchmark spanned four real systems (MOSTA, the axolotl telencephalon-regeneration atlas, *C. elegans* embryogenesis and zebrafish heart regeneration^33,35,37,38^) and two simulations with ground-truth lineage labels (expression-ambiguous but spatially separated lineages, and time-branching lineages, generated as Poisson counts over marker-block expression programs). It compared scBIOT-ST, non-spatial scBIOT, moscot-spatial and moscot^4^, Waddington-OT^9^ and Sinkhorn-OT^34^, each receiving the same expression representation. We reported top-1 transfer accuracy, normalized coupling entropy and recorded runtime. Normalized coupling entropy was the mean over source cells of each row-normalized coupling’s Shannon entropy divided by log *n*_target_ (range 0 to 1, lower indicating sharper couplings).

### Axolotl regeneration analysis

The same adjacent-stage strategy was applied to the ARTISTA axolotl telencephalon-regeneration Stereo-seq time course (2, 5, 10, 15, 20, 30 and 60 days post-injury)^35^, comprising 42,000 cells (6,000 subsampled per stage), 16,556 genes and 28 cell-type annotations. Counts were normalized to 10,000, log transformed and reduced to 2,000 highly variable genes; the L2-normalized autoencoder embedding was integrated across stages, and the forward field was computed with the same scale-matched spatial weighting (twice the embedding’s mean per-dimension standard deviation), 1,200 density-weighted centroids per stage, ordinal time and cell-type domains. Signed GTS used matched 6,000-cell subsamples per adjacent transition and the analytic Wald test with the same detection, false discovery rate and lineage-enrichment filters. Forward-fate composition and OT pullback traced the injury-induced progenitor cascade; controls comprised a shuffled-target pullback, a temporal-only coupling without the spatial term, and a background seed of random injured non-progenitor cells matched in number. Progenitor confinement was quantified as the injured-hemisphere cell fraction against this background, and the ependymoglia domain was scored by AUROC. All analyses used a fixed seed (0). These couplings describe coupling-derived ancestry and cell-state redistribution rather than validated clonal fate.

### Statistical analysis

Statistical tests were two-sided unless an explicitly directional gene-set hypothesis was declared. Independent sample groups used Mann-Whitney tests. Paired sequencing-sample effects used Wilcoxon signed-rank tests. Multiple testing used the Benjamini-Hochberg procedure within each declared family. Exact sample counts, effect definitions, tests and corrections are given in the relevant Methods and figure legends.

Analyses used Python (v3.12.8) with Scanpy (v1.12.1), AnnData (v0.12.16), NumPy (v2.4.4), pandas (v2.3.3), SciPy (v1.17.1), scikit-learn (v1.9.0), statsmodels (v0.14.6), scib-metrics (v0.5.9), POT (v0.9.6.post1), leidenalg (v0.11.0), Matplotlib (v3.10.9) and PyTorch (v2.10.0), together with the scBIOT package (v1.2.1). Mann-Whitney tests, Wilcoxon signed-rank tests, Spearman correlations and hypergeometric enrichment tests were performed using SciPy. Gene-level Benjamini-Hochberg false discovery rate correction for GTS was performed using statsmodels, and logistic regression with cross-validated AUROC was implemented using scikit-learn. The scVI and scANVI decoder-attribution baselines in Fig. 3 used scvi-tools v1.4.2. The exact software environment, command-line arguments, random seeds and analysis-specific scBIOT hyperparameters are documented in the archived environment and configuration files accompanying the versioned source code.

## Data availability

This study reused public datasets. HNSCC data are available in the Gene Expression Omnibus (GEO) under GSE139324 and GSE164690^28,29^. Paired multiome and CITE-seq BMMC data are available under GSE194122^31^. The Kang interferon-PBMC data are available in GEO under GSE96583^27^.

The Tabula Muris Senis data are available from Figshare (doi: 10.6084/m9.figshare.8273102.v2) and GEO under GSE132042^22^. Tabula Sapiens gene counts and metadata are available under GSE201333^23^. The MOSTA data are available from CNGB under CNP0001543^33^. Raw axolotl telencephalon-regeneration Stereo-seq data are available from CNGB under CNP0002068, and the processed ARTISTA resource used here is available from STOmicsDB under STDS0000056^35^. The *C. elegans* embryogenesis data are available under GSE126954^37^, and the zebrafish heart-regeneration data are available under GSE159032 and GSE158919^38^. Tahoe-100M is available from the Tahoe Bio dataset repository^30^.

The three Xenium spatial datasets (colorectal, liver and ovarian cancers) were obtained from the study of Ren et al.; raw sequencing data are available under HRA011129, image data under S-BIAD1900 and processed data through the SPATCH portal^24^. The multimodal PBMC reference used for label transfer is available under GSE164378^21^. Pancreas and lung benchmark objects were obtained from the scIB integration-task archive (doi: 10.6084/m9.figshare.12420968.v7)^1^. The two simulated spatiotemporal benchmark datasets were generated with the fixed seeds and parameters in source code. Source data and Extended Data, including processed numerical tables required to reproduce every plot, are deposited in the scBIOT repository.

## Code availability

The scBIOT source code and tutorials are available at https://github.com/haihuilab/scbiot and https://scbiot.readthedocs.io/en/stable/. The analyses used version 1.2.1. Figure-generation scripts, simulation parameters and input source tables are included in the same repository.

## Author contributions

H.Z. conceived the study, developed the method and software, performed the analyses, prepared the figures and wrote the original manuscript. P.Q. supervised the study, contributed resources and funding, and reviewed and edited the manuscript. Both authors interpreted the results and approved the final manuscript.

## Ethics, consent to participate, and consent to publish declarations

Not applicable.

## Acknowledgements

This work was supported in part by the National Natural Science Foundation of China (Grant No. 32350410397) and the Shenzhen Natural Science Fund (Grant No. 202205303001246).

## References

1. Luecken, M. D. et al. Benchmarking atlas-level data integration in single-cell genomics. Nature Methods 19, 41–50 (2022).

2. Argelaguet, R., Cuomo, A. S. E., Stegle, O. & Marioni, J. C. Computational principles and challenges in single-cell data integration. Nature Biotechnology 39, 1202–1215 (2021).

3. Heumos, L. et al. Best practices for single-cell analysis across modalities. Nature Reviews Genetics 24, 550–572 (2023).

4. Klein, D. et al. Mapping cells through time and space with moscot. Nature 638, 1065–1075 (2025).

5. Kang, J. B. et al. Efficient and precise single-cell reference atlas mapping with symphony. Nature Communications 12, 5890 (2021).

6. Xu, C. et al. Probabilistic harmonization and annotation of single-cell transcriptomics data with deep generative models. Molecular Systems Biology 17, e9620 (2021).

7. De Donno, C. et al. Population-level integration of single-cell datasets enables multi-scale analysis across samples. Nature Methods 20, 1683–1692 (2023).

8. Stuart, T. et al. Comprehensive integration of single-cell data. Cell 177, 1888–1902.e21 (2019).

9. Schiebinger, G. et al. Optimal-transport analysis of single-cell gene expression identifies developmental trajectories in reprogramming. Cell 176, 928–943.e22 (2019).

10. Forrow, A. & Schiebinger, G. LineageOT is a unified framework for lineage tracing and trajectory inference. Nature Communications 12, 4940 (2021).

11. Huguet, G. et al. Manifold interpolating optimal-transport flows for trajectory inference. Advances in neural information processing systems 35, 29705–29718 (2022).

12. Qu, R. et al. Gene trajectory inference for single-cell data by optimal transport metrics. Nature Biotechnology 43, 258–268 (2025).

13. Bunne, C., Schiebinger, G., Krause, A., Regev, A. & Cuturi, M. Optimal transport for single-cell and spatial omics. Nature Reviews Methods Primers 4, 58 (2024).

14. Cao, K., Gong, Q., Hong, Y. & Wan, L. A unified computational framework for single-cell data integration with optimal transport. Nature Communications 13, 7419 (2022).

15. Cang, Z. et al. Screening cell–cell communication in spatial transcriptomics via collective optimal transport. Nature Methods 20, 218–228 (2023).

16. Nitzan, M., Karaiskos, N., Friedman, N. & Rajewsky, N. Gene expression cartography. Nature 576, 132–137 (2019).

17. Haviv, D. et al. The covariance environment defines cellular niches for spatial inference. Nature Biotechnology 43, 269–280 (2025).

18. Demetci, P., Santorella, R., Sandstede, B., Noble, W. S. & Singh, R. SCOT: Single-cell multi-omics alignment with optimal transport. Journal of Computational Biology 29, 3–18 (2022).

19. Bunne, C. et al. Learning single-cell perturbation responses using neural optimal transport. Nature Methods 20, 1759–1768 (2023).

20. Dong, M. et al. Causal identification of single-cell experimental perturbation effects with CINEMA-OT. Nature Methods 20, 1769–1779 (2023).

21. Hao, Y. et al. Integrated analysis of multimodal single-cell data. Cell 184, 3573–3587.e29 (2021).

22. Tabula Muris Consortium. A single-cell transcriptomic atlas characterizes ageing tissues in the mouse. Nature 583, 590–595 (2020).

23. Tabula Sapiens Consortium. The tabula sapiens: A multiple-organ, single-cell transcriptomic atlas of humans. Science 376, eabl4896 (2022).

24. Ren, P. et al. Systematic benchmarking of high-throughput subcellular spatial transcriptomics platforms across human tumors. Nature Communications 16, 9232 (2025).

25. Brink, S. C. van den, Sage, F., Vergés-Ní, N. & Others. Single-cell sequencing reveals dissociation-induced gene expression in tissue subpopulations. Nature Methods 14, 935–936 (2017).

26. Lopez, R., Regier, J., Cole, M. B., Jordan, M. I. & Yosef, N. Deep generative modeling for single-cell transcriptomics. Nature Methods 15, 1053–1058 (2018).

27. Kang, H. M. et al. Multiplexed droplet single-cell RNA sequencing using natural genetic variation. Nature Biotechnology 36, 89–94 (2018).

28. Cillo, A. R. et al. Immune landscape of viral- and carcinogen-driven head and neck cancer. Immunity 52, 183–199.e9 (2020).

29. Kürten, C. H. L. et al. Investigating immune and non-immune cell interactions in head and neck tumors by single-cell RNA sequencing. Nature Communications 12, 7338 (2021).

30. Zhang, J. & Others. Tahoe-100M: A giga-scale single-cell perturbation atlas for context-dependent gene function and cellular modeling. bioRxiv 2025.02.20.639398 (2025) doi:10.1101/2025.02.20.639398.

31. Lance, C. et al. Multimodal single cell data integration challenge: Results and lessons learned. in Proceedings of machine learning research vol. 176 162–176 (2022).

32. Stoeckius, M. et al. Simultaneous epitope and transcriptome measurement in single cells. Nature Methods 14, 865–868 (2017).

33. Chen, A. et al. Spatiotemporal transcriptomic atlas of mouse organogenesis using DNA nanoball-patterned arrays. Cell 185, 1777–1792.e21 (2022).

34. Cuturi, M. Sinkhorn distances: Lightspeed computation of optimal transport. Advances in Neural Information Processing Systems 26, 2292–2300 (2013).

35. Wei, X. et al. Single-cell stereo-seq reveals induced progenitor cells involved in axolotl brain regeneration. Science 377, eabp9444 (2022).

36. Chizat, L., Peyré, G., Schmitzer, B. & Vialard, F.-X. Scaling algorithms for unbalanced optimal transport problems. Mathematics of Computation 87, 2563–2609 (2018).

37. Packer, J. S. et al. A lineage-resolved molecular atlas of C. elegans embryogenesis at single-cell resolution. Science 365, eaax1971 (2019).

38. Hu, B. et al. Origin and function of activated fibroblast states during zebrafish heart regeneration. Nature Genetics 54, 1227–1237 (2022).

