## Supplementary_File for "Informed, scalable and unified transport couplings for single-cell alignment and downstream inference with scBIOT"

**Extended Data**


Haihui Zhang[1,2*], Peiwu Qin[3]

[1]Department of Orofacial Sciences, School of Dentistry, University of California San Francisco, San Francisco, CA, USA
[2]Cancer Biology and Immunology Laboratory, University of California San Francisco, San Francisco, CA, USA
[3]Institute of Biopharmaceutical and Health Engineering, Shenzhen International Graduate School, Tsinghua University, Shenzhen, Guangdong, China

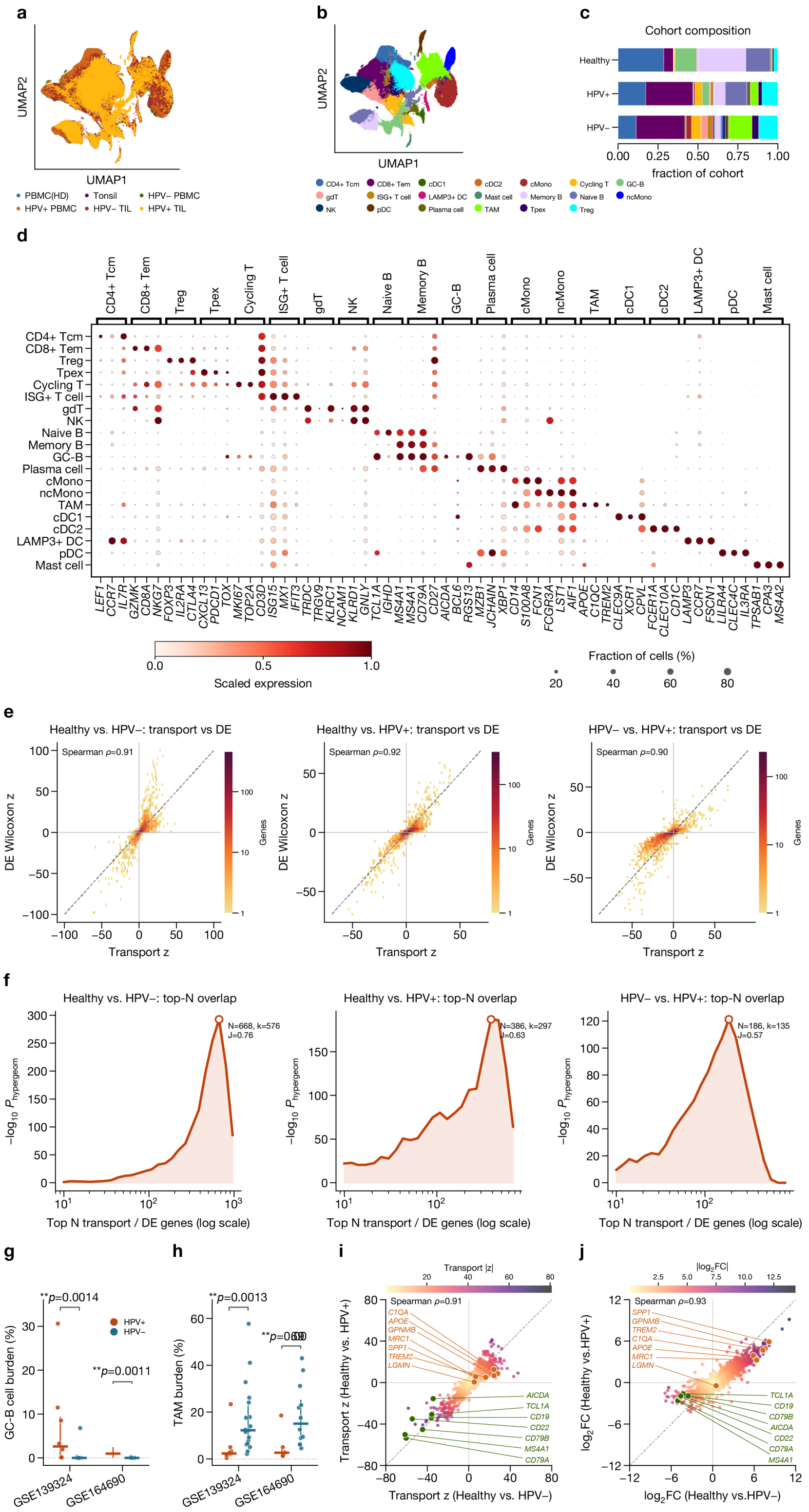
a
UMAP2
UMAP1
PBMC(HD)
Tonsil
HPV− PBMC
HPV+ PBMC
HPV− TIL
HPV+ TIL
b
UMAP2
UMAP1
CD4+ Tcm
CD8+ Tem
cDC1
cDC2
cMono
Cycling T
GC-B
gdT
ISG+ T cell
LAMP3+ DC
Mast cell
Memory B
Naive B
ncMono
NK
pDC
Plasma cell
TAM
Tpex
Treg
c
Cohort composition
Healthy
HPV+
HPV−
0.00
0.25
0.50
0.75
1.00
fraction of cohort
d
CD4+ Tcm
CD8+ Tem
Treg
Tpex
Cycling T
ISG+ T cell
gdT
NK
Naive B
Memory B
GC-B
Plasma cell
cMono
ncMono
TAM
cDC1
cDC2
LAMP3+ DC
pDC
Mast cell
LEF1 CCR7 IL7R GZMK CD8A NKG7 FOXP3 IL2RA CTLA4 CXCL13 PDCD1 TOX MKI67 TOP2A CD3D ISG15 MX1 IFIT3 TRDC TRGV9 KLRC1 NCAM1 KLRD1 GNLY TCL1A IGHD MS4A1 CD79A CD27 AICDA BCL6 RGS13 MZB1 JCHAIN XBP1 CD14 S100A8 FCN1 FCGR3A LST1 AIF1 APOE C1QC TREM2 CLEC9A XCR1 CPVL FCER1A CLEC10A CD1C LAMP3 CCR7 FSCN1 LILRA4 CLEC4C IL3RA TPSAB1 CPA3 MS4A2
0.0
0.5
1.0
Scaled expression
Fraction of cells (%)
20
40
60
80
e
Healthy vs. HPV−: transport vs DE
Spearman ρ=0.91
Healthy vs. HPV+: transport vs DE
Spearman ρ=0.92
HPV− vs. HPV+: transport vs DE
Spearman ρ=0.90
DE Wilcoxon z
Transport z
Genes
f
Healthy vs. HPV−: top-N overlap
N=668, k=576
J=0.76
Healthy vs. HPV+: top-N overlap
N=386, k=297
J=0.63
HPV− vs. HPV+: top-N overlap
N=186, k=135
J=0.57
−log10 P hypergeom
Top N transport / DE genes (log scale)
g
**p=0.0014
**p=0.0011
HPV+
HPV−
GC-B cell burden (%)
GSE139324
GSE164690
h
**p=0.0013
**p=0.039
TAM burden (%)
GSE139324
GSE164690
i
Transport |z|
Spearman ρ=0.91
C1QA
APOE
GPNMB
MRC1
SPP1
TREM2
LGMN
AICDA
TCL1A
CD19
CD22
CD79B
MS4A1
CD79A
Transport z (Healthy vs. HPV+)
Transport z (Healthy vs. HPV−)
j
|log2FC|
Spearman ρ=0.93
SPP1
GPNMB
TREM2
C1QA
APOE
MRC1
LGMN
TCL1A
CD19
CD79B
AICDA
CD22
CD79A
MS4A1
log2FC (Healthy vs.HPV+)
log2FC (Healthy vs.HPV−)

**Extended Data Fig. 1 | Annotation and robustness of the integrated head and neck squamous cell carcinoma (HNSCC) immune atlas. a,b,** scBIOT-integrated CD45-positive atlas colored by tissue/human papillomavirus (HPV) cohort (**a**) and curated immune-cell identity (**b**). The tissue-restricted analysis contains 124,506 cells from 49 samples (5 healthy tonsils, 14 HPV-positive tumors and 30 HPV-negative tumors). **c,** Immune-cell composition in each cohort. **d,** Expression of canonical markers used to support the cell-state annotations; dot size denotes the fraction of expressing cells and color denotes scaled mean expression. **e,** Signed gene transport score (GTS) $z$ score versus Wilcoxon differential-expression $z$ score for healthy versus HPV-negative, healthy versus HPV-positive and HPV-negative versus HPV-positive contrasts. Spearman $\rho$ measures rank concordance; cell-level scores localize effects but do not constitute biological replicates. **f,** Hypergeometric enrichment of overlap among genes ranked by transport and differential expression relative to the analyzed gene universe. **g,h,** Germinal-center B (GC-B) cell (**g**) and macrophage (**h**) burden in HPV-positive and HPV-negative tumors, shown separately for GSE139324 and GSE164690. Each point is one sample; zero-count samples are retained. Brackets show two-sided Mann-Whitney tests within each source study. The within-study panels test direction and robustness rather than providing an independent validation cohort. **i,j,** Shared tumor-associated and HPV-associated transport and expression effects for the leading macrophage (TAM) and germinal-center/tertiary lymphoid structure (GC/TLS) B-cell genes, the same two gene blocks shown in Fig. 4i. TAM genes are elevated and GC/TLS B-cell genes reduced in both healthy-versus-tumor contrasts. The same direction held within GSE139324 alone (all five healthy tonsils and 26 tumor samples), removing study-of-origin confounding; because healthy tissue came from only this study, external replication remains absent. The TAM genes are restricted to those enriched in the annotated TAM compartment over both classical and non-classical monocytes; the plasma/Ig/UPR control block is not highlighted.

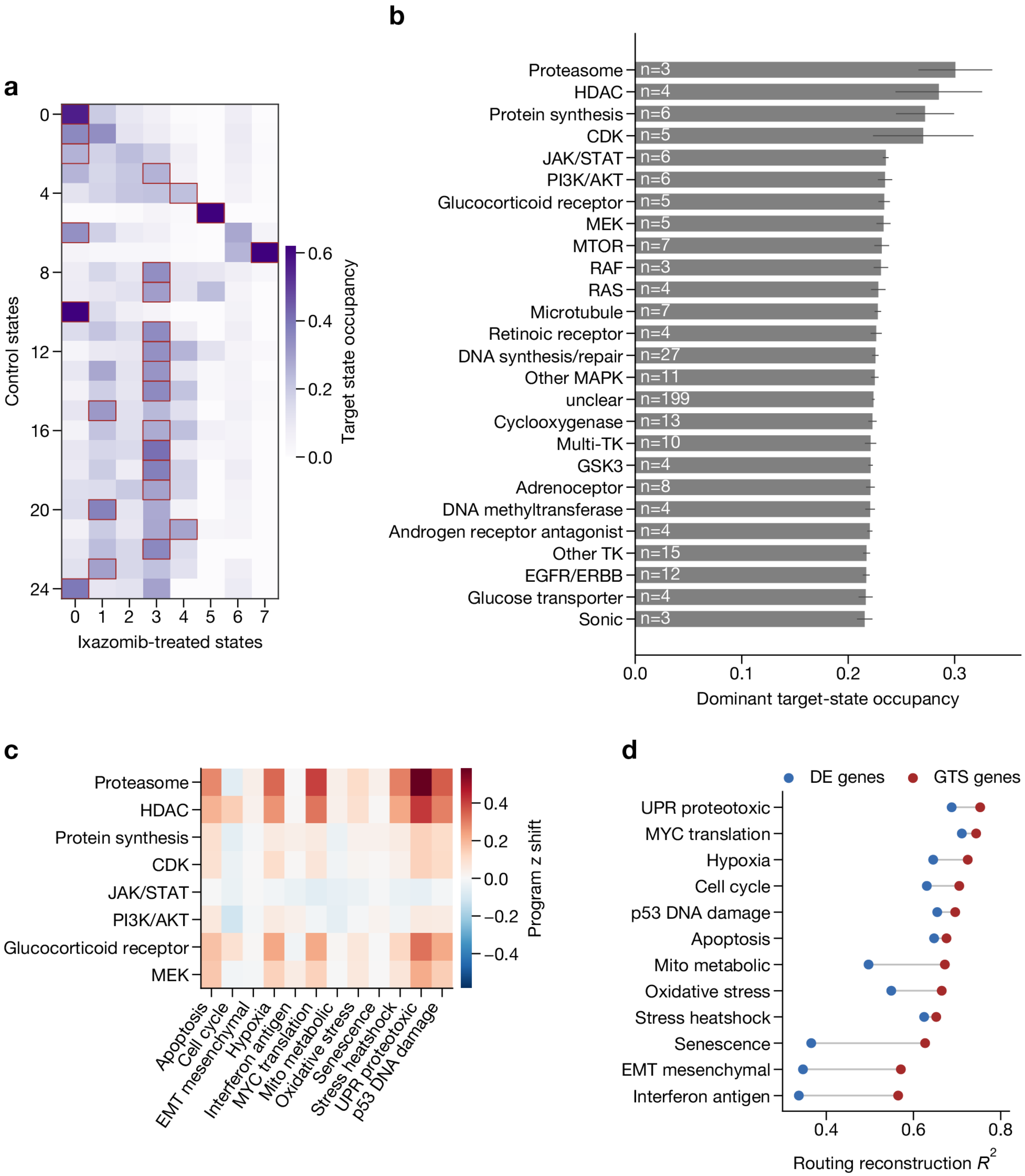
a
Control states
Ixazomib-treated states
Target state occupancy
0.6
0.4
0.2
0.0
b
Proteasome n=3
HDAC n=4
Protein synthesis n=6
CDK n=5
JAK/STAT n=6
PI3K/AKT n=6
Glucocorticoid receptor n=5
MEK n=5
MTOR n=7
RAF n=3
RAS n=4
Microtubule n=7
Retinoic receptor n=4
DNA synthesis/repair n=27
Other MAPK n=11
unclear n=199
Cyclooxygenase n=13
Multi-TK n=10
GSK3 n=4
Adrenoceptor n=8
DNA methyltransferase n=4
Androgen receptor antagonist n=4
Other TK n=15
EGFR/ERBB n=12
Glucose transporter n=4
Sonic n=3
Dominant target-state occupancy
c
Proteasome
HDAC
Protein synthesis
CDK
JAK/STAT
PI3K/AKT
Glucocorticoid receptor
MEK
Apoptosis
Cell cycle
EMT mesenchymal
Hypoxia
Interferon antigen
MYC translation
Mito metabolic
Oxidative stress
Senescence
Stress heatshock
UPR proteotoxic
p53 DNA damage
Program z shift
d
DE genes
GTS genes
UPR proteotoxic
MYC translation
Hypoxia
Cell cycle
p53 DNA damage
Apoptosis
Mito metabolic
Oxidative stress
Stress heatshock
Senescence
EMT mesenchymal
Interferon antigen
Routing reconstruction $R^2$

**Extended Data Fig. 2 | Prototype transport retains structured state routing in Tahoe-100M.** The analysis used 95.6 million cells retained after filtering, spanning 379 drugs, 50 cancer cell lines and 26 mechanism-of-action groups, including an unannotated "unclear" catch-all group. **a,** Row-normalized source-to-target prototype occupancy for a representative ixazomib perturbation. Rows sum to one; boxes identify the target state with the largest occupancy for each source state. **b,** Dominant target-state occupancy summarized by mechanism group. Drugs are the replication unit; bar height is the mean across drugs and error bars denote the standard error of the mean (s.e.m.) across drugs. **c,** For the eight mechanism groups with the highest dominant target-state occupancy (the leading groups in **b**), the median target-minus-source shift across gene programs after target states were weighted by coupling occupancy. **d,** Reconstruction of the observed source-to-target routing for each response program using genes ranked by marginal differential expression (DE genes) or coupling-specific GTS (GTS genes). Across programs, GTS-ranked genes reconstructed the routing at least as well as differentially expressed genes. This comparison assesses routing reconstruction and does not establish biological information beyond differential expression.

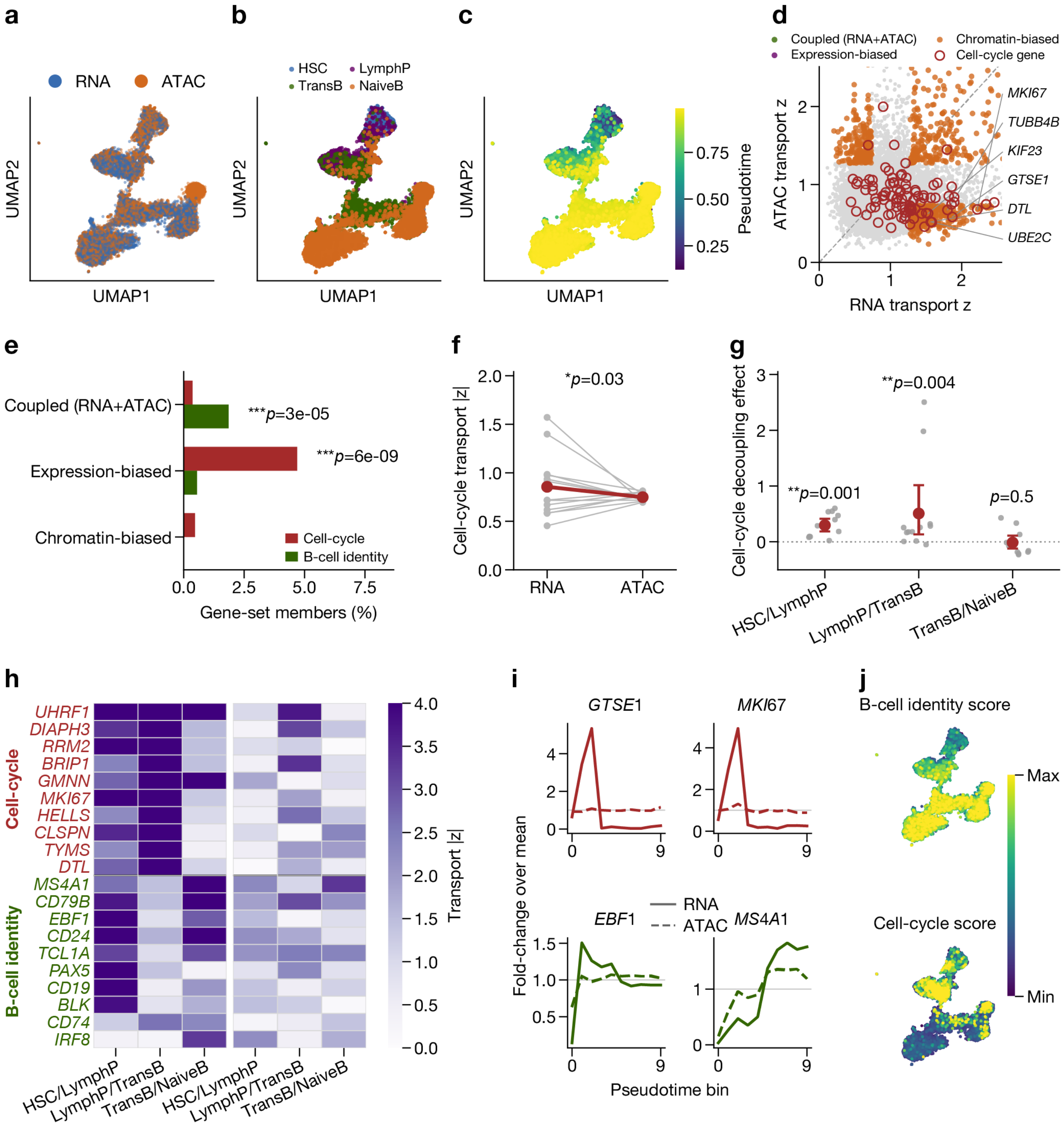
a
RNA
ATAC
UMAP2
UMAP1
b
HSC
LymphP
TransB
NaiveB
UMAP2
UMAP1
c
Pseudotime
0.75
0.50
0.25
UMAP2
UMAP1
d
Coupled (RNA+ATAC)
Expression-biased
Chromatin-biased
Cell-cycle gene
ATAC transport z
RNA transport z
MKI67
TUBB4B
KIF23
GTSE1
DTL
UBE2C
e
Coupled (RNA+ATAC)
Expression-biased
Chromatin-biased
***p=3e-05
***p=6e-09
Cell-cycle
B-cell identity
Gene-set members (%)
f
Cell-cycle transport |z|
*p=0.03
RNA
ATAC
g
Cell-cycle decoupling effect
**p=0.001
**p=0.004
p=0.5
HSC/LymphP
LymphP/TransB
TransB/NaiveB
h
Cell-cycle
UHRF1
DIAPH3
RRM2
BRIP1
GMNN
MKI67
HELLS
CLSPN
TYMS
DTL
B-cell identity
MS4A1
CD79B
EBF1
CD24
TCL1A
PAX5
CD19
BLK
CD74
IRF8
Transport |z|
HSC/LymphP
LymphP/TransB
TransB/NaiveB
HSC/LymphP
LymphP/TransB
TransB/NaiveB
i
GTSE1
MKI67
EBF1
MS4A1
RNA
ATAC
Fold-change over mean
Pseudotime bin
j
B-cell identity score
Cell-cycle score
Max
Min

**Extended Data Fig. 3 | A separate B-lymphoid analysis reproduces expression-biased cell-cycle transport.** **a-c,** Joint representation of 10,713 paired multiome cells colored by RNA or assay for transposase-accessible chromatin (ATAC) view (**a**), B-lineage stage (**b**) and pseudotime (**c**). **d,** Dynamic RNA and ATAC transport magnitudes for shared genes; dashed thresholds define coupled, expression-biased, chromatin-biased and untransported classes. **e,** Fractions of cell-cycle and B-cell-identity genes in each declared transport class. **f,** Whole-lineage cell-cycle transport in RNA and ATAC across 13 sequencing samples. Lines connect measurements from the same sample; the two-sided Wilcoxon signed-rank $P$ value is 0.033. **g,** Baseline-adjusted RNA-minus-ATAC effects for individual B-lineage transitions, with Benjamini-Hochberg (BH) correction across transitions. **h-j,** Step-specific transport heat maps (**h**), representative gene trends along pseudotime (**i**) and cell-cycle or B-cell-identity program scores (**j**).

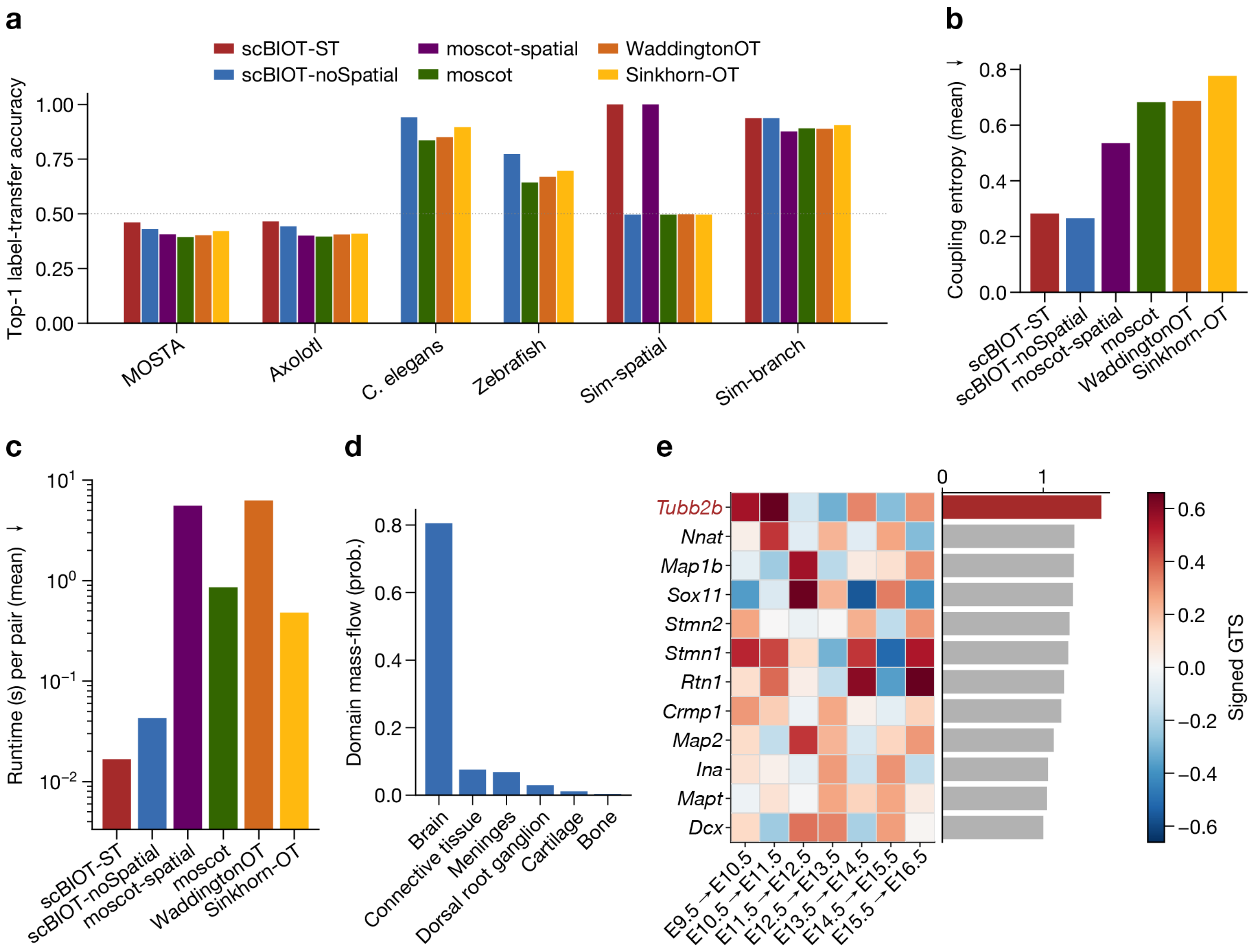
a
scBIOT-ST
scBIOT-noSpatial
moscot-spatial
moscot
WaddingtonOT
Sinkhorn-OT
Top-1 label-transfer accuracy
MOSTA
Axolotl
C. elegans
Zebrafish
Sim-spatial
Sim-branch
b
Coupling entropy (mean)
c
Runtime (s) per pair (mean)
d
Domain mass-flow (prob.)
Brain
Connective tissue
Meninges
Dorsal root ganglion
Cartilage
Bone
e
Tubb2b
Nnat
Map1b
Sox11
Stmn2
Stmn1
Rtn1
Crmp1
Map2
Ina
Mapt
Dcx
E9.5→E10.5
E10.5→E11.5
E11.5→E12.5
E12.5→E13.5
E13.5→E14.5
E14.5→E15.5
E15.5→E16.5
Signed GTS

**Extended Data Fig. 4 | Spatiotemporal benchmark and neural-domain transport.** **a,** Top-1 domain-transfer accuracy for scBIOT-ST, non-spatial scBIOT, moscot-spatial, moscot, Waddington-OT and Sinkhorn-OT across four real and two simulated datasets. Every method received the same expression representation; bars are descriptive dataset-level point estimates. **b,** Mean normalized coupling entropy across the same tasks; lower values indicate more concentrated, not necessarily more accurate, couplings. **c,** Mean recorded runtime per adjacent-stage pair. Runtime depends on implementation, hardware and dataset size and was not subjected to a population-level test. **d,** Target-domain mass for a selected Spinal cord-to-Brain coupling in the Mouse Organogenesis Spatiotemporal Transcriptomic Atlas (MOSTA). **e,** Lineage-restricted signed GTS for the neural-domain flow after detection, false discovery rate (FDR) and lineage-enrichment filtering; *Tubb2b* is highlighted. OT, optimal transport.

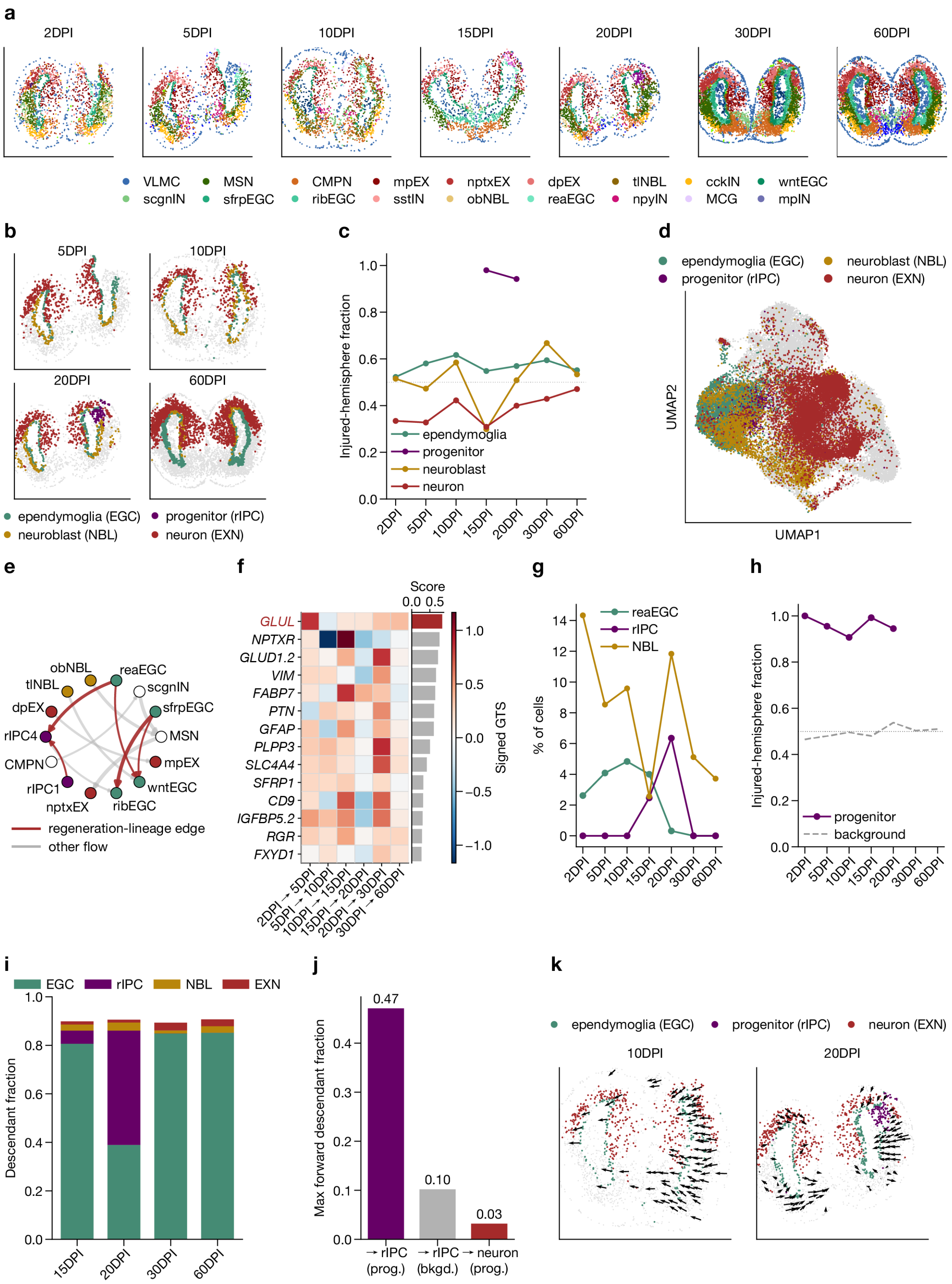
a
2DPI
5DPI
10DPI
15DPI
20DPI
30DPI
60DPI
VLMC
MSN
CMPN
mpEX
nptxEX
dpEX
tINBL
cckIN
wntEGC
scgnIN
sfrpEGC
ribEGC
sstIN
obNBL
reaEGC
npyIN
MCG
mpIN
b
ependymoglia (EGC)
progenitor (rIPC)
neuroblast (NBL)
neuron (EXN)
c
Injured-hemisphere fraction
d
UMAP1
UMAP2
e
regeneration-lineage edge
other flow
f
Score
Signed GTS
GLUL
NPTXR
GLUD1.2
VIM
FABP7
PTN
GFAP
PLPP3
SLC4A4
SFRP1
CD9
IGFBP5.2
RGR
FXYD1
g
% of cells
h
Injured-hemisphere fraction
progenitor
background
i
Descendant fraction
EGC
rIPC
NBL
EXN
j
Max forward descendant fraction
0.47
0.10
0.03
→ rIPC (prog.)
→ rIPC (bkgd.)
→ neuron (prog.)
k
ependymoglia (EGC)
progenitor (rIPC)
neuron (EXN)
10DPI
20DPI

**Extended Data Fig. 5 | Spatiotemporal transport recovers the injury-induced progenitor cascade of axolotl telencephalon regeneration.** Adjacent-stage scBIOT-ST couplings were fitted across an axolotl brain-regeneration Stereo-seq time course (2 to 60 days post-injury, DPI)[1]. **a,** Telencephalon sections across DPI colored by cell type. **b,** Regeneration-lineage cells localize to the injured hemisphere at representative stages. **c,** Injured-hemisphere fraction of each lineage across DPI; regeneration intermediate progenitors (rIPC) are strongly biased to the injured hemisphere. **d,** scBIOT latent space colored by regeneration lineage using uniform manifold approximation and projection (UMAP). **e,** Directed cell-state OT flow network; regeneration-lineage edges tracing the reactive-ependymoglia-to-progenitor cascade are highlighted. **f,** Signed GTS across adjacent-DPI transitions for the top-ranked regeneration-associated genes, with per-gene score bars; *GLUL* is highlighted. **g,** Progenitor-wave cell fractions across DPI, showing reactive ependymoglia (reaEGC) early followed by rIPC and neuroblasts (NBL). **h,** Injured-hemisphere fraction of progenitors versus a background control across DPI. **i,** Forward-transport target composition at 15 to 60 DPI. **j,** Maximum forward-transport fraction for progenitor-to-rIPC transport relative to a background control and to progenitor-to-neuron transport; the recovered cascade stops short of the rare terminal progenitor-to-neuron transport. **k,** Projected spatial velocity field in the regenerating wound at 10 and 20 DPI. The inferred couplings describe a spatially wound-confined, temporally ordered progenitor cascade and provide coupling-derived correspondences rather than validated clonal fate.

1. Wei, X. *et al.* Single-cell stereo-seq reveals induced progenitor cells involved in axolotl brain regeneration. *Science* **377**, eabp9444 (2022).
